# NF1 Loss Remodels Tumor Niches for Immune Evasion

**DOI:** 10.64898/2026.01.11.698818

**Authors:** Milad Ibrahim, Irineu Illa-Bochaca, Tara Muijlwijk, Ines Delclaux, Katherine S Ventre, George Jour, Paola Angulo Salgado, Shi Qiu, Agrima Dutt, Amanda W. Lund, Iman Osman, Markus Schober

## Abstract

Genetic and transcriptional alterations in cancer cells shape their interactions with immune and stromal compartments, influencing tumor progression, immune evasion, and response to immune checkpoint inhibitors (ICI). Yet, how these interactions are organized within tumor architecture and linked to clinical outcomes remains unclear. Neurofibromin 1 (*NF1*) is a tumor suppressor gene that is frequently inactivated across multiple cancer types. *NF1* loss-of-function mutations occur in up to 27% of melanoma cases and are associated with poor clinical outcomes. Here, we used spatial multi-omics analysis to uncover 12 meta-niches, each comprising distinctive cell types with distinct characteristics, in human melanoma tissues. We found that niches containing immunosuppressive cancer-associated fibroblasts (CAFs) and macrophages were significantly enriched in *NF1* mutant melanoma (*NF1^Mut^*) tissues. In contrast, niches containing cytotoxic CD8 T cells were significantly diminished. *NF1* loss correlates with increased epidermal growth factor signaling (EGFR) signaling and reduced antigen presentation in tissues with limited CD8 T cell infiltration in both human and mouse melanoma. We demonstrate that EGFR inhibition restores antigen presentation and activates immune responses in a syngeneic *Nf1* knockdown model resistant to ICIs. These data, therefore, define functionally distinctive niches enriched in *NF1^Mut^* melanoma that likely contribute to their aggressive nature and nominate EGFR signaling as a specific target to reinvigorate ICI responses. We therefore link an understudied genetic driver to specific immune architectures and ultimately therapy resistance and suggest a therapeutic strategy expected to improve treatment outcomes in *NF1^Mut^* melanoma patients.

## Introduction

Genetic driver mutations have been linked to cancer-defining histopathological features,^1^ immune evasion,^2^ and therapeutic responses.^3^ Transcriptional heterogeneity, which may emerge as a function of environmental interactions, may further shape histopathological heterogeneity even in tumors driven by the same mutations.^4,5^ Still, how the interaction between tumor cells and their environment, which drives these histopathology-defining niches, selects for specific mechanisms of immune evasion, treatment resistance, and clinical outcomes in patients remains poorly understood.

Cutaneous melanoma is the deadliest type of skin cancer due to its high metastatic potential and resistance to treatment.^6,7^ Melanoma is genetically, transcriptionally, and histopathologically heterogeneous.^1–3,8^ Most melanomas are driven by oncogenic mutations in *BRAF* or *NRAS*, or by loss-of-function mutations in the tumor suppressor gene *Neurofibromin* 1 (*NF1*).^7^ *NF1* is mutated in up to 27% of melanomas and in up to 45% of *BRAF-* and *NRAS-wild*-type tumors.^9^ Despite being associated with worse disease-specific and overall survival,^10^ *NF1* mutant (*NF1^Mut^*) melanomas lack effective targeted therapy.^9,11,12^ Moreover, co-occurring *NF1* mutations with *BRAF* or *NRAS* mutations, confer resistance to BRAF and MEK inhibitors.^9,11–13^ As a result, clinical management of *NF1^Mut^* melanoma remains particularly challenging.

*NF1^Mut^* melanomas have the highest tumor mutational burden, and NF1 loss has been linked to increased programmed death-ligand 1 (PD-L1) cell surface expression, suggesting a heightened likelihood of response to immune checkpoint inhibitors (ICI).^3,10,11,14,15^ Paradoxically, more than half of *NF1^Mut^* melanomas fail to respond to ICI.^9,15^ We have previously shown that *NF1^Mut^*melanomas are more aggressive due to distinct transcriptional programs characterized by increased proliferation driven by receptor tyrosine kinases (RTKs), particularly epidermal growth factor receptor (EGFR).^9,11^ Although EGFR signaling contributes to immune evasion and ICI resistance in EGFR-driven head and neck squamous cell carcinomas^16,17^ and lung cancers^18^, it is unclear whether this pathway plays similar roles in melanoma, specifically in the context of *NF1^Mut^*melanomas resistant to ICI. We postulate that elucidating mechanisms of immune evasion enriched in *NF1^Mut^* patients is essential for developing more effective therapeutic strategies tailored specifically for *NF1^Mut^*patients, who have no other available targeted therapies.

Here, we have leveraged spatial multi-omic approaches to characterize cellular neighborhoods comprising melanoma cells in various transcriptionally distinct cell states and functionally distinct CAFs, endothelial cells, myeloid cells, T cells, and other immune and stromal cell types in human *NF1^Mut^* and *NF1* wild-type (*NF1^WT^*) melanoma tissues. We have defined cellular neighborhoods observed across multiple melanoma tissues as meta-niches (MNs), and shown that their abundance, composition, and functions are governed by NF1-dependent gene expression programs. We have identified EGFR-dependent immune evasion mechanisms in patient tissues and in a preclinical *Nf1* knockdown melanoma model that is resistant to PD-1 inhibition but becomes responsive after EGFR inhibition.

## Results

### NF1-loss impacts meta-niche composition and architecture in melanoma

We analyzed spatial transcriptomic and proteomic profiles of melanoma tissues at the single-cell level **(Figure 1A, Table S1)** using tissue microarrays comprising 34 *NF1^Mut^* and 30 *NF1^WT^* tissue cores. 42 of these cores were profiled using the 10X Genomics 5K Xenium platform. After data acquisition, cell segmentation, and rigorous quality control, we removed cells with fewer than 50 transcripts and tissues with fewer than 5000 cells. We batch-corrected and normalized the data across independent tissues to create an integrated data set of 1,003,604 high-quality cells from 39 patients (20 *NF1^Mut^*and 19 *NF1^WT^*). Leiden clustering distinguished melanoma cells from B cells, T cells, myeloid cells, plasma cells, epithelial cells, and other stromal cell types **(Figure 1B)**. We annotated the cell types based on their expression of lineage-defining marker genes **(Figure 1C).** Although these cell types were detected in most melanoma tissues, their distribution varied considerably **(Figure 1D).** The heterogeneous spatial distribution of melanoma and stromal cells suggests that each tumor tissue contains multiple specialized niches, and the organization of these cell populations may dictate tumor behavior, immune interactions, and histopathological features.

**Figure 1.**
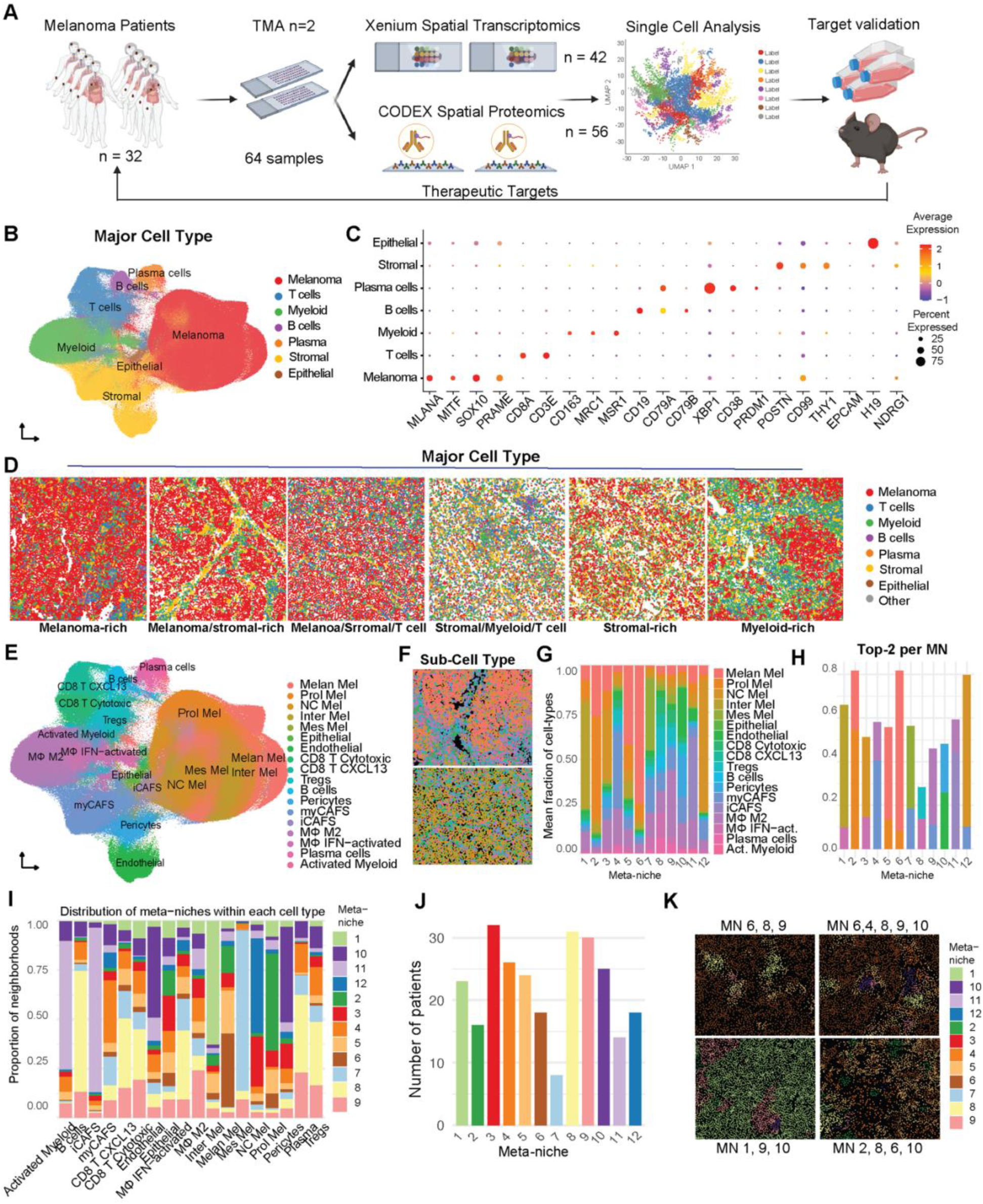
Distinct spatial niches and cellular neighborhood architectures of melanoma. **(A)** Schematic representation of the study workflow, starting from isolating 64 melanoma tumors from 32 patients, generating TMAs, performing Xenium spatial transcriptomics on 42 samples and CODEX spatial proteomics on 56 samples to perform single-cell analysis, and testing targets identified in preclinical models. **(B)** Uniform manifold approximation and projection (UMAP) plot of major cell types identified in the Xenium dataset. **(C)** Dot plot showing canonical marker genes for each cell type identified in (B). **(D)** Representative field of view (FOV) of a Xenium image, showing major cell types in multiple melanoma samples. **(E)** UMAP plot of lineage sub-types identified in the Xenium dataset. **(F)** Representative FOV of a Xenium image showing lineage sub-types identified in (E). **(G, H)** Stacked-bar plot showing the composition of each meta-niche across all cell types (G) or top 2 cell types (H). **(I)** Stacked-bar plot showing the abundance of each cell type across all meta-niches. **(J)** Bar plot showing the number of patients included in each meta-niche. **(K)** Representative FOV images of meta-niches identified in melanoma patients.

To better understand the biological relevance of this observation, we re-clustered cells from each lineage based on their gene expression profiles into 18 distinct cell types, including five melanoma cell states **(Figure 1E-F, S1A).** We identified melanoma cells with melanocytic, proliferative, intermediate, neural crest (NC)-like, and mesenchymal (MES) features consistent with previous reports.^19^ Immune cells included cytotoxic, CXCL13-positive CD8 T cells, regulatory T cells (Tregs), M2-like and interferon (IFN)-stimulated macrophages, activated myeloid cells, B cells, and plasma cells. Stromal cells could be subclassified into cancer-associated fibroblasts (CAFs) with myofibroblast-like (myCAF) and inflammatory (iCAF) features, pericytes, endothelial cells, and epithelial cell types.

To define complex cellular neighborhoods in *NF1^Mut^* and *NF1^WT^* melanoma tissues, we mapped their cell-type identities and transcriptional programs back to each tissue core. We retrieved the centroid of each cell in the tissue and analyzed its cellular neighborhood within a 40 µM radius of each index cell using a fast radius-based nearest-neighbor search (frNN).^20^ Cells with fewer than three neighboring cells (omitting the index cell) were excluded from further analyses. Next, we used k-means clustering^21^ to group these neighborhoods into 12 frequently occurring meta-niches (MNs), which we define here as cellular spatial neighborhood patterns that occur in several independent tissue samples **(Figure 1G)**. We classified these MNs based on the two most frequently occurring cell types within each MN **(Figure 1H)** and found that each cell type was present in multiple niches **(Figure 1I)**. Each MN was observed in multiple samples from different patients **(Figure 1J**), with multiple MNs per sample **(Figure 1K, S1B)**, indicating that these patterns are consistent across the cohort rather than patient-specific.

Comparing *NF1^Mut^* and *NF1^WT^* revealed that their tumors contain the same major cell types **(Figure S1C-D)**. However, *NF1^Mut^* melanoma tumors were significantly enriched for MN3, comprising neural crest (NC)-like melanoma cells and immune-suppressive M2-like macrophages, and for MN12, comprising NC-like melanoma cells and immune-suppressive myCAFs **(Figure 2A-B, S2A)**. Notably, the NC-like melanoma state has been associated with ICI resistance.^22^ Conversely, MN10, which is enriched in pericytes and endothelial cells, was less commonly observed in *NF1^Mut^* melanoma tissues **(Figure 2A, C)**, even though the overall population sizes of pericytes and endothelial cells in *NF1^Mut^*melanoma tissues were similar to those of *NF1^WT^* tumors **(Figure S2B)**. These data suggest that *NF1^WT^* melanoma tissues contain more mature vascular structures than *NF1^Mut^* melanoma tissues, which have been associated with improved tumor-immune responses.^23^ Consistent with this idea, MN8, which is enriched in cytotoxic T cells, Tregs, and IFN-activated macrophages, was less frequently detected in *NF1^Mut^* melanoma tissues. These data suggest that *NF1^Mut^* melanoma employs different immune evasion strategies and is less infiltrated by activated immune cells.

**Figure 2.**
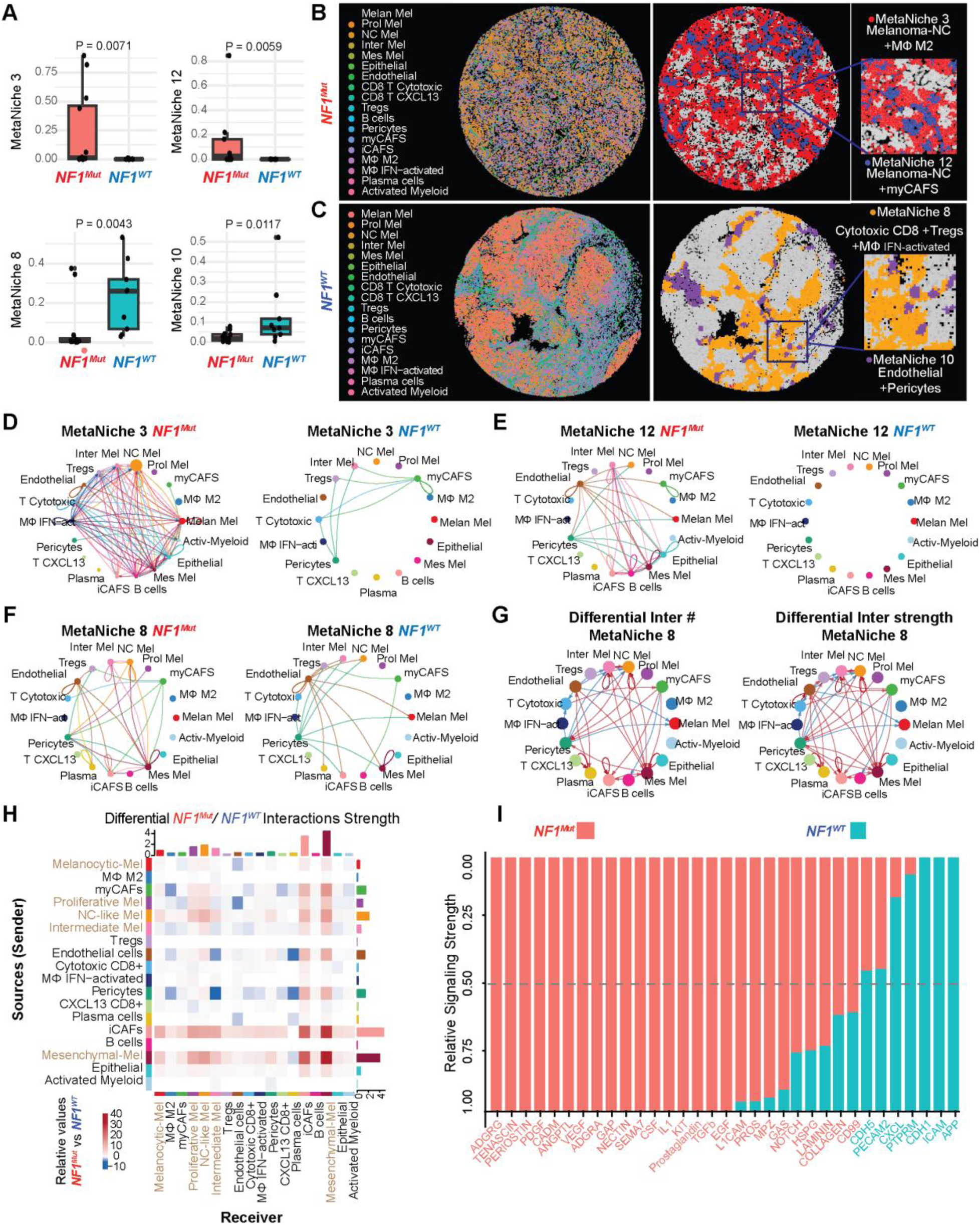
*NF1* mutant melanomas are enriched in oncogenic and immune suppressive meta-niches. **(A)** Bar plots and data points of spatial niches in *NF1^Mut^* or *NF^WT^* melanoma. P-values indicate a Wilcox test. **(B, C)** Representative field of view (FOV) of Xenium images, showing cell types and niches that are either increased in *NF1^Mut^* (B) or *NF^WT^* (C) melanoma. **(D-G)** CellChat circle plots showing signaling patterns in Meta-niche 3 for *NF1^Mut^*or *NF^WT^* (D), signaling patterns in Meta-niche 12 for *NF1^Mut^*or *NF^WT^* (E), signaling patterns in Meta-niche 8 for *NF1^Mut^* or *NF^WT^* (F), or differential number of interactions or differential interaction strength *NF1^Mut^*or *NF^WT^* meta-niche 8 (G). Red arrows indicate more and stronger interactions, while blue arrows indicate fewer or weaker interactions. **(H)** CellChat heatmaps showing *NF1^Mut^* vs. *NF^WT^* differential interactions strength. Senders are in rows and receivers are in columns. Red color indicates stronger, while blue color indicates weaker interactions in *NF1^Mut^*. **(I)** CellChat Bar plot showing cell-cell signaling in *NF1^Mut^* Vs. *NF^WT^* microenvironment. Red label indicates signaling significantly higher in *NF1^Mut^*, teal label indicates signaling significantly higher in *NF^WT^,* and black label indicates no significant difference.

We then probed our data set for mechanisms of cell-cell communication and adhesion within each niche and tested how NF1 loss influenced these interactions using CellChat.^24^ MN3, which is more abundant in *NF1^Mut^* melanoma **(Figure 2B)**, is dominated by signals from NC-like melanoma cells to melanoma cells in all other states, as well as to stromal and immune cells that characterize MN3 **(Figure 2D)**. The most characteristic and abundant signaling mechanisms in MN3 include Myelin Protein Zero (MPZ) and L1 Cell Adhesion Molecule (L1CAM), which are commonly associated with tumor progression, invasion, and metastasis **(Figure S2C)**.^25^ MN12 is also more abundant in *NF1^Mut^* melanoma tissue, but it is dominated by interactions between NC-like melanoma cells and myCAFs, with fewer interactions between tumor and immune cells **(Figure 2E).**

MN8 was dominated by immune cell interactions and contained activated macrophages signaling to melanoma cells, with stronger signaling in *NF1^WT^* than in *NF1^Mut^*melanoma tissues **(Figure 2F)**. Differential analysis of cell-cell signaling patterns in MN8 further revealed more receptor-ligand interactions among cytotoxic CD8 T cells, regulatory T cells, and activated macrophages in *NF1^WT^* melanoma tissues (**Figure 2G**). Conversely, MN8 showed more interactions between melanoma cells and CAFs in *NF1^Mut^*melanoma tissues. These signaling patterns result in significantly increased CXCL signaling in *NF1^WT^* MNs. In contrast, Platelet Endothelial Cell Adhesion Molecule (PECAM2), MPZ, Cell Adhesion Molecule (CADM), Protease-Activated Receptors (PARs), Heparan Sulfate Proteoglycans (HSPG), and Thrombospondin (THBS) signaling were significantly increased in MNs of *NF1^Mut^* melanoma samples (**Figure S2D)**, indicating a highly angiogenic, immune-suppressed tumor microenvironment (TME).

To investigate global signaling dynamics and compare them with signaling within each MN, we performed CellChat across all cell types identified in our cohort. Differences in interaction number and strength between *NF1^Mut^* and *NF1^WT^*melanoma indicated that cells within MNs engage in more robust communication. For example, NC-like melanoma cells transmitted more and stronger signals to myCAFs and M2-like macrophages **(Figure 2H, S2E)**. Intriguingly, myCAFs sent collagen and laminin signals back to support NC-like cells in *NF1^Mut^*melanoma, indicating crosstalk between these two cell types. Conversely, myCAFs interacted less with cytotoxic CD8 T cells and CXCL13 CD8 T cells in *NF1^Mut^* melanoma tissues. Collagen, laminin, epidermal growth factor (EGF), transforming growth factor (TGF) beta, KIT proto-oncogene-receptor tyrosine kinase (KIT), prostaglandin, vascular endothelial growth factor (VEGF), platelet-derived growth factor (PDGF), and L1CAM signaling were significantly increased in the *NF1^Mut^* TME, while CXCL signaling was downregulated **(Figure 2I)**. Collagen signaling was also increased in NC-like cells and myCAFs in *NF1^Mut^* melanoma tissues compared with *NF1^WT^* melanoma **(Figure S2F, G)**, indicating that *NF1^Mut^* melanoma could be more fibrotic. Moreover, myCAFs and iCAFs exhibited increased signaling for PDGF, EGF, TGFβ, KIT, VEGF, adhesion G protein-coupled receptors (ADGR) A and D, tenascin, and laminin in *NF1^Mut^* melanoma. Collectively, these data suggest that *NF1^Mut^* melanoma is characterized by a more fibrotic microenvironment, increased EGF signaling, fewer intact blood vessels, and reduced immune signaling.

### NF1 mutant tumors accumulate dysfunctional CD8 T cells

The MNs we uncovered in *NF1^Mut^* and *NF1^WT^* melanomas suggested that they could reflect global differences in the spatial organization between melanoma cells and their TME. To test this, we measured the distance between *NF1^Mut^* or *NF1^WT^* melanoma cells and other cell types in their TME. This global neighborhood analysis showed that *NF1^Mut^* melanoma cells tend to be closer to CAFs, Tregs, M2-like macrophages, and pericytes than *NF1^WT^* melanoma cells. In contrast, they were farther away from cytotoxic CD8 T cells, CXCL13 CD8 T cells, and endothelial cells in *NF1^Mut^* melanoma tissues **(Figure 3A-B)**. These results indicate that the TME surrounding *NF1^Mut^* melanoma cells is depleted of immune cells but enriched in CAFs, whereas CD8 T cells and other immune cell types infiltrate *NF1^WT^*melanoma tissues.

**Figure 3.**
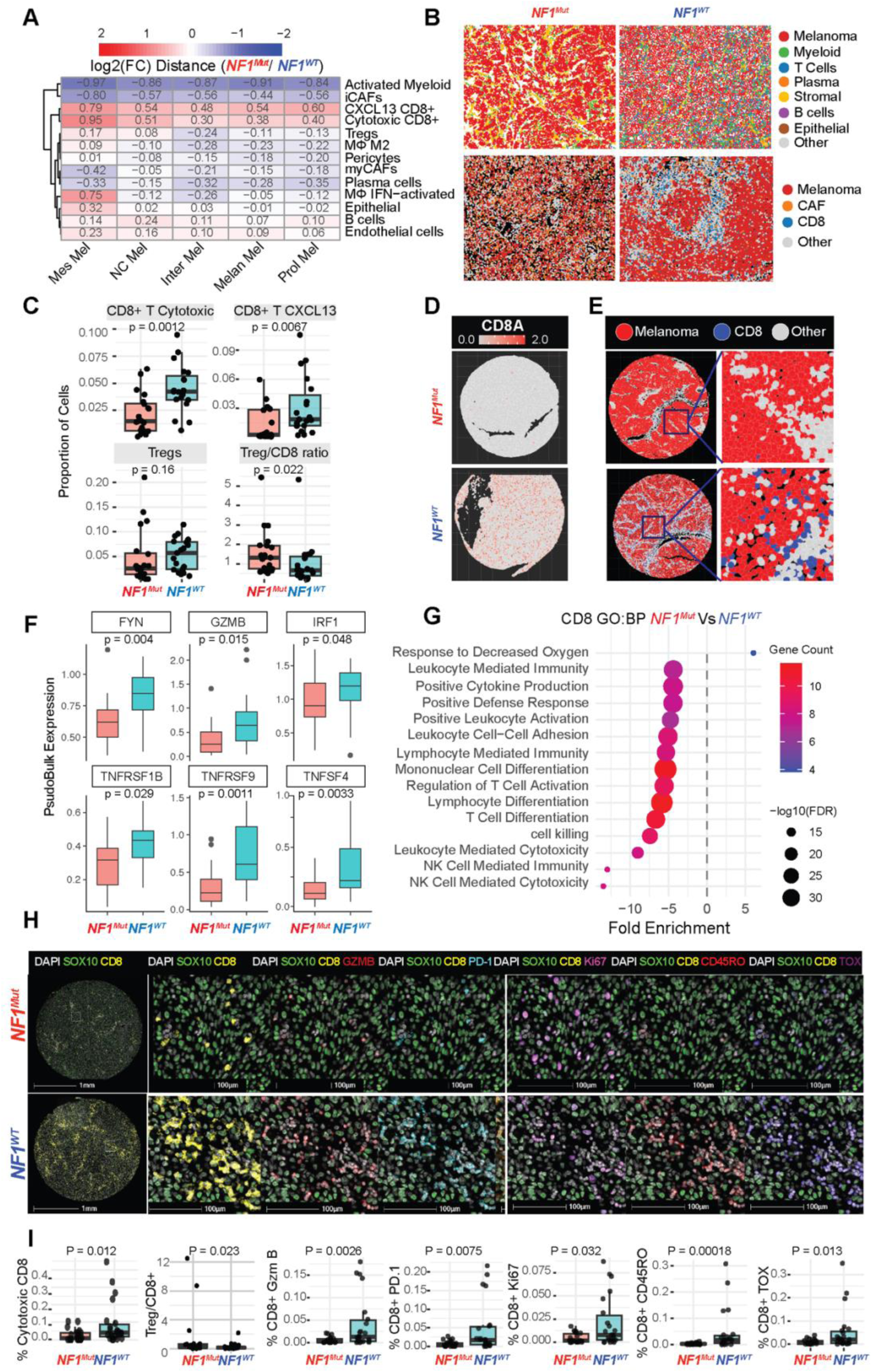
*NF1* mutant melanoma is characterized by decreased T cell infiltration and activity. **(A)** *NF1^Mut^* Vs. *NF^WT^* differential heatmap showing the distance between different states of melanoma cells in columns and microenvironment cell types in rows. Red color indicates cells are farther away, while blue color indicates the cells are closer to *NF1^Mut^* tumors. **(B)** Representative field of view (FOV) images showing the spatial organization of tumor, CD8, and stromal cells, shown by major or sub-cell type in *NF1^Mut^* or *NF^WT^*. **(C)** Box plots and data points of CD8 T, CXCL13 CD8 T, and Tregs over the total number of cells per patient or the Treg/CD8 ratio. P-values indicate a Wilcox test. **(D, E)** Spatial heatmap (D) or FOV Images (E) of CD8 T cells in *NF1^Mut^*or *NF^WT^* melanoma patients. **(F)** Box plots showing differentially expressed genes comparing *NF1^Mut^* to *NF^WT^* melanoma CD8 T cell clusters using pseudo bulk. P-values indicate a Wilcox test. **(G)** GO pathway analysis shows that response to decreased oxygen is upregulated, while T cell activation, differentiation, and NK activity are downregulated in CD8 T cell clusters of *NF1^Mut^* melanoma. **(H)** Representative images showing tumor-infiltrating CD8 T cells/total number of cells per patient and the co-localization of CD8 activity markers GZMB, PD-1, Ki67, CD45 RO, and TOX. **(I)** Box plots and data points of CD8 activity markers are shown in (H). P-values indicate a Wilcox test.

*NF1^Mut^* melanoma cells were surrounded by fewer cytotoxic CD8 T cells and CXCL13 CD8 T cells, and the ratio of regulatory T cells to CD8 T cells was higher in these tumors, consistent with a suppressed immune TME **(Figure 3C-E)**. Fewer CD8 T cells infiltrated *NF1^Mut^* tumors, and they expressed significantly less *FYN, GZMB, IRF1, TNFRSF1B, TNFSF4,* and *TNFRSF9* **(Figure 3F).** These changes were enriched for pathways associated with decreased cytokine production, lower T cell activation and differentiation, and reduced T cell-mediated killing of cancer cells (**Figure 3G).**

To cross-validate our Xenium results using an independent approach, we examined the same cohort with a custom 45-plex antibody panel. We imaged 54 tissues using the Akoya-PhenoCycler platform, and used HALO software to quantify, segment, and annotate 1,079,175 single cells in these tissues. This approach captured melanoma cells, as well as CAFs, endothelial cells, dendritic cells (DC), macrophages, CD4 T cells, CD8 T cells, memory T cells, NK cells, and B cells. We retained 1,052,420 cells from 49 samples after quality control. These immunostainings confirmed that *NF1^Mut^* melanoma tissues contained fewer cytotoxic CD8 T cells and a higher regulatory T cell-to-CD8 T cell ratio. Furthermore, markers of cytotoxic (GZMB), memory (CD45RO), exhausted (PD-1, TOX), and activated/proliferating (Ki67) CD8 T cells were all significantly decreased in *NF1^Mut^* melanoma tissues (**Figure 3H, I**). CD4/CD8/DC intratumoral immune triads have recently been shown to be required for ICI responses.^26^ We quantified the presence of these triads using proximity scores based on distances between CD4, CD8, and DCs (CD11c+, HLA.DR+, CD14-, CD68-, SOX10-). We observed significantly fewer CD4/CD8/DC immune complexes in *NF1^Mut^* melanoma tissues compared to *NF1^WT^* melanoma tissues **(Figure S3A-B).**

Consistent with these results, imputation of gene expression data of skin cutaneous melanoma (SKCM) that were generated by the Cancer Genome Atlas Network (TCGA)^14^ with CIBERSORTx,^27^ using single-cell RNA sequencing data from Tirosh et al.^8^ as a reference, also detected significantly fewer CD8 T cells in *NF1^Mut^*melanoma samples than in *NF1^WT^* melanomas (**Figure S3B**).

### NF1 loss results in decreased melanoma antigen presentation

Our analyses uncovered distinct immune evasion strategies in *NF1^WT^*and *NF1^Mut^* melanomas. *NF1^WT^* tumors were enriched for immune-active MNs in which CD8 T cells followed a classical activation, memory formation, and exhaustion trajectory, whereas *NF1^Mut^*melanomas were dominated by immune-silenced MNs with broadly attenuated CD8 T cell states and reduced tumor immune interactions. Consistent with this, *NF1^Mut^* immune TME exhibited significantly lower expression scores of inflammatory cytokines (*IL1B, IL6, IL12A/B, IL17A, IL18, TNF, IFNG, IFNA1,* and *IFNB1*), chemokines (*CXCL9, CXCL10, CXCL11, and CXCL13*), and chemokine receptors (*CCR5, CCR7, CXCR3, CXCR4,* and *CXCR5*) within the *NF1^Mut^* immune microenvironment **(Figure 4A)**. These changes also correlate with suppressed immune stimulatory (*CD80, CD86, ICOS, CD28,* and *TNFRSF9*) and elevated immune inhibitory checkpoint (*PDCD1, CD274, CTLA4, LAG3, HAVCR2, TIGIT,* and *VSIR*) gene expression scores. Together, these findings suggest that *NF1^Mut^* melanomas are less effectively recognized by immune cells, pointing to impaired antigen presentation rather than dominant checkpoint-mediated inhibition. Cancer cells frequently reduce or lose expression of Human Leukocyte Antigen (HLA) as a primary immune evasion strategy.^28^ Our highly multiplexed IHC analyses detected significantly less HLA expression in *NF1^Mut^*melanoma cells **(Figure 4B-C).** This reduced HLA expression correlates directly (r_s_ = 0.8355) with fewer CD8 T cells in tumor tissues **(Figure 4D)**. We also identified a lower expression of *TAP1, TAP2, CANX,* and *CIITA* in *NF1^Mut^* melanoma cells than in *NF1^WT^* melanoma cells in our Xenium data **(Figure 4E)**.

**Figure 4.**
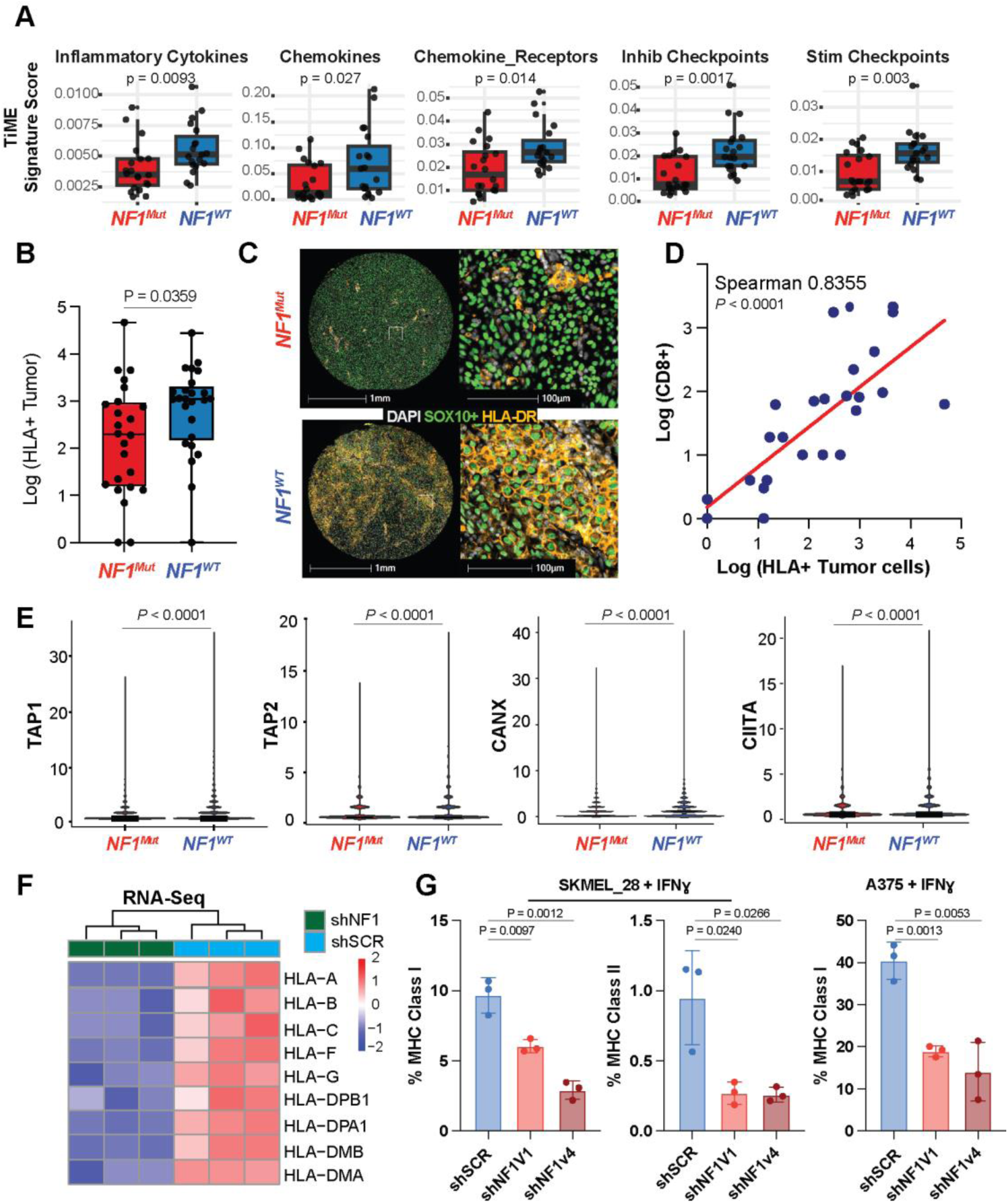
*NF1* loss inhibits MHC class I and II antigen presentation. **(A)** Box plots and data points showing immune signature gene scores identified in the immune microenvironment of *NF1^Mut^* and *NF^WT,^* normalized to the total number of cells per patient. P-values indicate a Wilcox test. **(B)** Box plots and data points of log (HLA+ tumor cells). P-values indicate a Wilcox test. **(C)** Representative images showing melanoma tumor cells stained with HLA+ in *NF1^Mut^* and *NF^WT^*tumor cells. **(D)** Scatter plot showing correlation and regression analysis of the number of CD8 T cells and the number of HLA+ tumor cells. P-values indicate Spearman correlation. **(E)** Violin plots of the expression of MHC-related genes in melanoma cells measured by spatial transcriptomics in *NF1^Mut^* and *NF^WT^* melanoma cells. P-values indicate a Wilcox test. **(F)** Heatmap showing differentially expressed MHC class I and II genes in A375 melanoma cells following *NF1* knockdown. **(G)** Flow cytometry bar graphs and data points of MHC class I or II after interferon stimulation with (shNF1) or without (shSCR) NF1 knockdown in SKMEL-28 or A375 melanoma cell lines.

To test if NF1 loss was directly responsible for the reduced expression of HLAs in *NF1^Mut^* melanoma, we transduced A375 melanoma cells with non-targeting control short-hairpin RNAs (shSCR) or *NF1*-inhibiting short-hairpin RNAs (shNF1). Differential gene expression analyses revealed significantly lower *HLA-A, HLA-B, HLA-C, HLA−F, HLA−G, HLA−DMA, HLA−DMB, HLA−DPA1,* and *HLA−DPB1* expression in shNF1 transduced A375 cells compared to shSCR controls **(Figure 4F)**. Furthermore, *NF1*-knockdown A375 cells expressed less MHC-I, and SK-MEL28 cells expressed less MHC-I and MHC-II on their cell surface compared to shSCR-expressing control cells, even after IFNɣ-stimulation (**Figure 4G)**. These data demonstrate that NF1 loss reduces HLA expression, and IFNɣ-stimulation is unable to fully restore it.

### EGF-EGFR signaling is enriched in *NF1*-mutant melanoma meta-niches and mediates immune evasion

Based on our previously reported data showing that NF1 loss activates EGFR signaling, which is essential for tumor proliferation and survival,^9,11^ we sought to investigate the role of EGFR in *NF1^Mut^* melanoma MNs, immune evasion, and the associated loss of HLA expression in *NF1^Mut^* melanoma cells. To do this, we isolated melanoma, stromal, and myeloid cell types to identify gene sets significantly enriched or depleted in *NF1^Mut^* or *NF1^WT^* melanoma tissues. Transcriptional programs enriched in *NF1^Mut^* melanoma cells include EMT, UV response, hypoxia, TGFβ, and Tumor Necrosis Factor alpha (TNFα) signaling **(Figure S4A)**. These pathways were driven by significantly increased expression of *NOTCH1, SOX2, NGFR, ERBB2, EGFR, TGFB2, TGBR2, VEGFA, VEGFC,* and *PDGFRB* **(Figure S4B)**. Conversely, *NF1^WT^* melanoma cells were enriched for pathways associated with increased inflammatory responses, IFNα and IFNɣ responses, and IL2-STAT5 and IL6-STAT3 signaling, driven by significantly higher expression of IL10, IL21, IFNG, IFNL1, IFNA17, and TNFSF18.

Furthermore, CAFs we identified in *NF1^Mut^* melanoma tissues upregulated genes involved in cell division, cell-cycle regulation, and EGF signaling. They also downregulated genes related to antigen processing, antigen presentation, cytokine production, inflammatory response, T cell activation, and adaptive immune responses **(Figure S4C)**. Similarly, myeloid cells we identified in *NF1^Mut^* melanoma tissues expressed transcripts associated with elevated growth factor responsiveness, increased TGFβ receptor signaling, reduced inflammatory response, dampened immune effector processes, and diminished cytokine and immune responses **(Figure S4D)**. Notably, EGF signaling was elevated in CAFs, myeloid cells, and *NF1^Mut^*melanoma cells. These data suggest that EGFR signaling could promote oncogenic and immune-evasive mechanisms in *NF1^Mut^* melanoma.

Next, we identified gene sets that were up- or downregulated in the MNs identified in this study. Differential gene expression analyses revealed that gene sets associated with hypoxia, TGFβ, and epithelial-to-mesenchymal transition (EMT) were consistently upregulated, whereas IFNɣ and inflammatory response genes were consistently downregulated in MNs of *NF1^Mut^*melanoma **(Figure 5A)**. To determine whether and how EGFR signaling affects immune evasion and tumor growth in *NF1^Mut^* melanoma, we examined EGFR expression across all MNs in our *NF1^Mut^* and *NF1^WT^* melanoma tissue collection. EGFR expression was higher and more prevalent in *NF1^Mut^*tumors across most MNs **(Figure 5B, C)**. To explore the biological programs associated with elevated EGFR in *NF1^Mut^*tumors, we assessed pathway activities and their associations with EGFR expression in melanoma cells within each MN. Differential pathway analysis revealed that increased EGFR expression in *NF1^Mut^* tumors was associated with increased EMT, hypoxia, and TGFβ signaling, as well as decreased inflammation, IFN responses, IL6–JAK–STAT3 signaling, IL2–STAT5 signaling, and p53 signaling **(Figure 5D)**. These data demonstrate that elevated EGFR expression correlates with activation of oncogenic and stress-response programs and suppression of immune-related transcriptional programs in *NF1^Mut^* melanoma tissues. EGF signaling was observed between melanoma cells and CAFs in MNs of *NF1^Mut^* melanoma but not *NF1^WT^* melanoma **(Figure S2F-G).**

**Figure 5.**
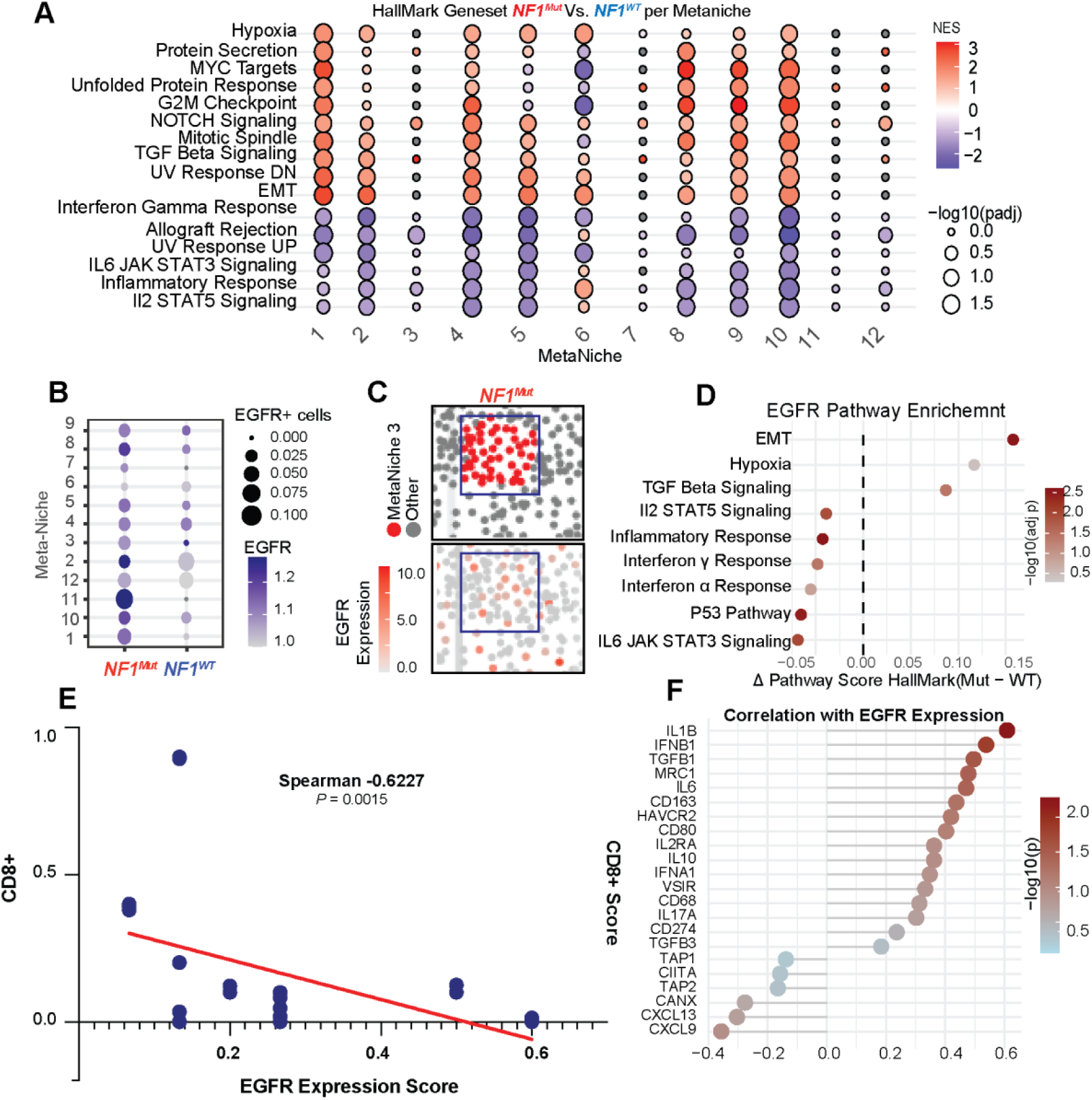
EGF-EGFR signaling promotes an immune evasive microenvironment in *NF1* mutant melanoma. **(A)** Dot plot showing differential transcriptional Hallmark gene sets enriched in each meta niche comparing *NF1^Mut^* to *NF^WT^* melanoma. Color represents fold change, and size represents adjusted p-value. **(B)** Dot plot showing EGFR expression in each meta-niche, comparing *NF1^Mut^* and *NF^WT^* tumors. The size of the dots represents the fraction of EGFR-positive cells, while the color intensity reflects the mean EGFR expression. **(C)** Representative images showing MN3 and EGFR expression in *NF1^Mut^* microenvironment. **(D)** Differential pathway activity in melanoma cells associated with EGFR, calculated as the mean difference in pathway module scores between *NF1^Mut^* and *NF^WT^*. Color represents the statistical significance. **(E)** Scatter plot showing correlation and regression analysis of the number of CD8 T cells and the EGFR immunostaining expression score. P-values indicate Spearman correlation. **(F)** Spearman correlation analysis of EGFR expression with immune regulatory genes in melanoma clusters of *NF1^Mut^* tissue.

EGFR expression is negatively correlated (r_s_ = -0.622) with CD8 T cell abundance in *NF1^Mut^* melanoma tissues, indicating that tumors with the highest EGFR expression have the fewest CD8 T cells **(Figure 5E)**. Increased EGFR expression also correlates with reduced expression of antigen presentation genes, including *TAP1, TAP2, CANX,* and *CIITA,* together with decreased levels of the immune-stimulatory chemokines CXCL13 and CXCL9 **(Figure 5F).**^29,30^ Conversely, *EGFR* expression is directly associated with higher expression of *TGFB1, TGFB3, IL1B, IL10, CD274,* and *HAVCR2,* all of which can enhance immune evasion in *NF1^Mut^* melanoma by inhibiting antigen presentation, decreasing T cell proliferation, and activating immune checkpoints.

To assess which of the observed immune-evasive phenotypes can be attributed to NF1 loss, we compared differentially expressed genes between shNF1 knockdown and shSCR expressing A375 cells. These analyses linked gene sets associated with increased angiogenesis, hypoxia, EMT, TGF-β, and NOTCH signaling (**Figure 6A**) as well as increased expression of *VEGFA, ERBB2, EGFR, VEGFC, TGFβR2,* and *NOTCH1* (**Figure 6B**) to NF1 loss. Conversely, ectopic expression of a doxycycline (dox)-inducible recombinant NF1 (rNF1) transgene (**Figure 6C**) inhibited gene sets associated with these pathways (**Figures 6D-E**). Consistent with these gene expression changes, we found that *NF1^Mut^* melanoma cells grew more slowly upon re-expressing rNF1 **(Figure 6F)**. These results suggest that NF1 inhibits EGFR, whereas NF1 loss activates EGFR, to promote cell proliferation and immune evasion.

**Figure 6.**
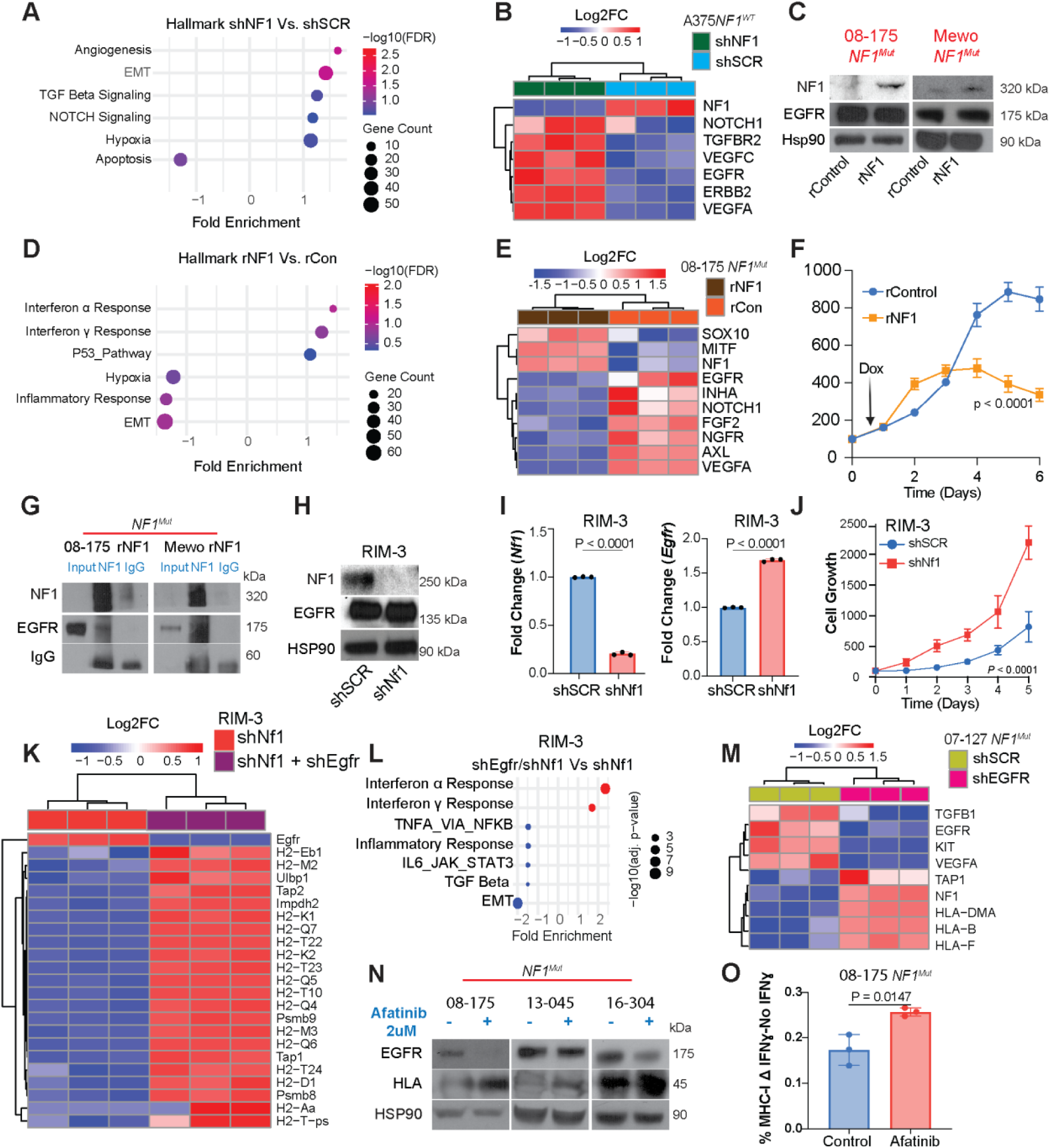
EGFR inhibition restores HLA antigen presentation. **(A)** GSEA Hallmark pathway analysis showing EMT, TGFβ, angiogenesis, and hypoxia are upregulated as a result of *NF1* knockdown **(B).** Heatmap showing differentially expressed genes between shNF1 and shSCR expressing cells. **(C)** Western blots showing NF1 and EGFR expression before and after DOX-induced NF1 expression (rNF1) compared to control (rControl) in *NF1^Mut^* STC-08-175 and Mewo melanoma cells. HSP90 served as a loading control. **(D)** GSEA Hallmark pathway analysis showing IFNα and IFNɣ responses are upregulated, and EMT, hypoxia and inflammatory response are downregulated as a result of NF1 ectopic expression. **(E)** Heatmap showing significant differential expressed genes comparing NF1 ectopic expression to control. **(F)** Growth curves of *NF1^Mut^* STC-08-175 melanoma cells ectopically expressing rNF1 compared to control. *P*-values were calculated with a two-sided t-test at the experimental endpoint. Data points represent mean ± SD. (n=12). **(G)** Western blots of NF1 co-immunoprecipitation showing EGFR levels after pulling down with NF1 or IgG control antibodies after DOX-induced NF1 expression (rpbNF1) or control (rbpControl) in STC-08-175 or Mewo *NF1^Mut^* melanoma cells. **(H)** Western blots comparing NF1 and EGFR, expression in shNf1 compared to shSCR RIM-3 murine melanoma cells. HSP90 served as a loading control. **(I)** RT-PCR data showing *Nf1* and *Egfr* expression changes in shNf1compared to shSCR expressing RIM3 melanoma cells. Bar graphs indicate mean ± SD (n=3). *P*-values were calculated with a two-sided t-test. **(J)** Growth curves of RIM-3 melanoma cells expressing shNf1 compared to shSCR (control). *P*-values were calculated with a two-sided t-test at the experimental endpoint. Data points represent mean ± SD. (n=12). **(K)** Heatmap showing differentially expressed antigen presentation genes in RIM-3-shNF1 melanoma cells after EGFR knockdown. **(L)** GSEA Hallmark Gene Sets analysis showing enriched pathways in shEGFR RIM-3-shNF1 cell lines. **(M)** Heatmap showing differentially expressed HLA genes in shEGFR and shSCR (control) expressing *NF1^Mut^* STC-07-127 melanoma cells. **(N)** Western blots comparing HLA and EGFR expression with and without afatinib treatment in three human *NF1^Mut^* melanoma cell lines. HSP90 served as a loading control. **(O)** Flow cytometry analysis showing MHC class I antigen presentation difference between IFNɣ-stimulated or non-stimulated *NF1^Mut^* STC-08-175 melanoma cells with and without afatinib treatment. Bar graphs indicate mean ± SD (n=3). *P*-values were calculated with a two-sided t-test.

BioGrid network data showed that NF1 can physically interact with EGFR.^31^ Therefore, we co-immunoprecipitated NF1 and EGFR from cell lysates of *NF1^Mut^* 08-175 or Mewo cells expressing rNF1, confirming that NF1 can physically interact with EGFR in melanoma cells **(Figure 6G)**, and their interaction may inhibit the activation of EGFR, thus initiating its transcriptional suppression by interfering with a positive feedback circuit that drives the expression of EGFR in *NF1^Mut^* melanoma cells.

The increased expression of EGFR in *NF1^Mut^* melanomas and its association with immune evasion mechanisms suggest that EGFR inhibition could reactivate tumor immunity and thereby improve the response to ICI therapy. To test this hypothesis, we used syngeneic RIM-3 cells,^32^ which were isolated from a melanoma that developed in *Tyr::Nras^Q61K^ Cdkn2a^Fl/Fl^* mice.^33^ sh*Nf1* knockdown decreased NF1 expression and increased Egfr expression in RIM3 cells (**Figure 6H-I**), significantly accelerating their growth rate (**Figure 6J)**. These results in RIM3 cells are consistent with data we recently reported on human melanoma short-term cultures and xenograft models,^11^ and could now allow us to functionally test how NF1 loss and/or EGFR activation affect immune evasion and ICI response

To do this, we first knocked down *Egfr* in shNf1-expressing RIM3 cells and used RNA-seq to identify gene sets regulated by EGFR. We found a significant decrease in *Egfr* expression, along with increased expression of genes required for antigen presentation and NK and T cell activation (**Figure 6K**). Pathway analyses also revealed increased IFNα and IFNɣ responses, while TNFα signaling via NFκB, EMT, and TGFβ signaling were downregulated as a result of EGFR inhibition (**Figure 6L**). We observed similar changes in HLA expression when we compared shEGFR to shSCR expressing human *NF1^Mut^* melanoma cell lines **(Figure 6M**) or when we treated human *NF1^Mut^* melanoma patients derived short-term cultures (STCs) with the EGFR inhibitor afatinib **(Figure 6N).**^34^ Furthermore, MHC-I expression increased even more when we treated *NF1^Mut^*melanoma cells with afatinib and IFNɣ than when treating them with IFNɣ alone **(Figure 6O)**.

These data demonstrate that increased EGFR signaling compromises antigen presentation in *NF1^Mut^*melanoma cells, and these changes could contribute to immune evasion, defective T cell infiltration, cytotoxic tetrad formation, and ICI resistance.

### EGFR inhibition induces an immune response that synergizes with immune checkpoint inhibitors in *NF1*-depleted melanoma mouse models

To assess whether NF1 loss affects immune evasion, we orthotopically transplanted equal numbers of RIM3 cells expressing shSCR or shNf1 into the dermis of immunocompromised *Nude* or syngeneic C57BL/6J mice and assessed differences in tumor initiation rates. Although both shSCR and shNf1-expressing RIM3 cells initiated tumors at similar rates in *Nude* mice **(Figure 7A)**, only shNF1-expressing RIM3 cells were able to form tumors in C57BL/6J mice (**Figure 7B**). This result supports the notion that NF1 loss enhances immune evasion, consistent with our spatial transcriptomic results from human melanoma tissues.

**Figure 7.**
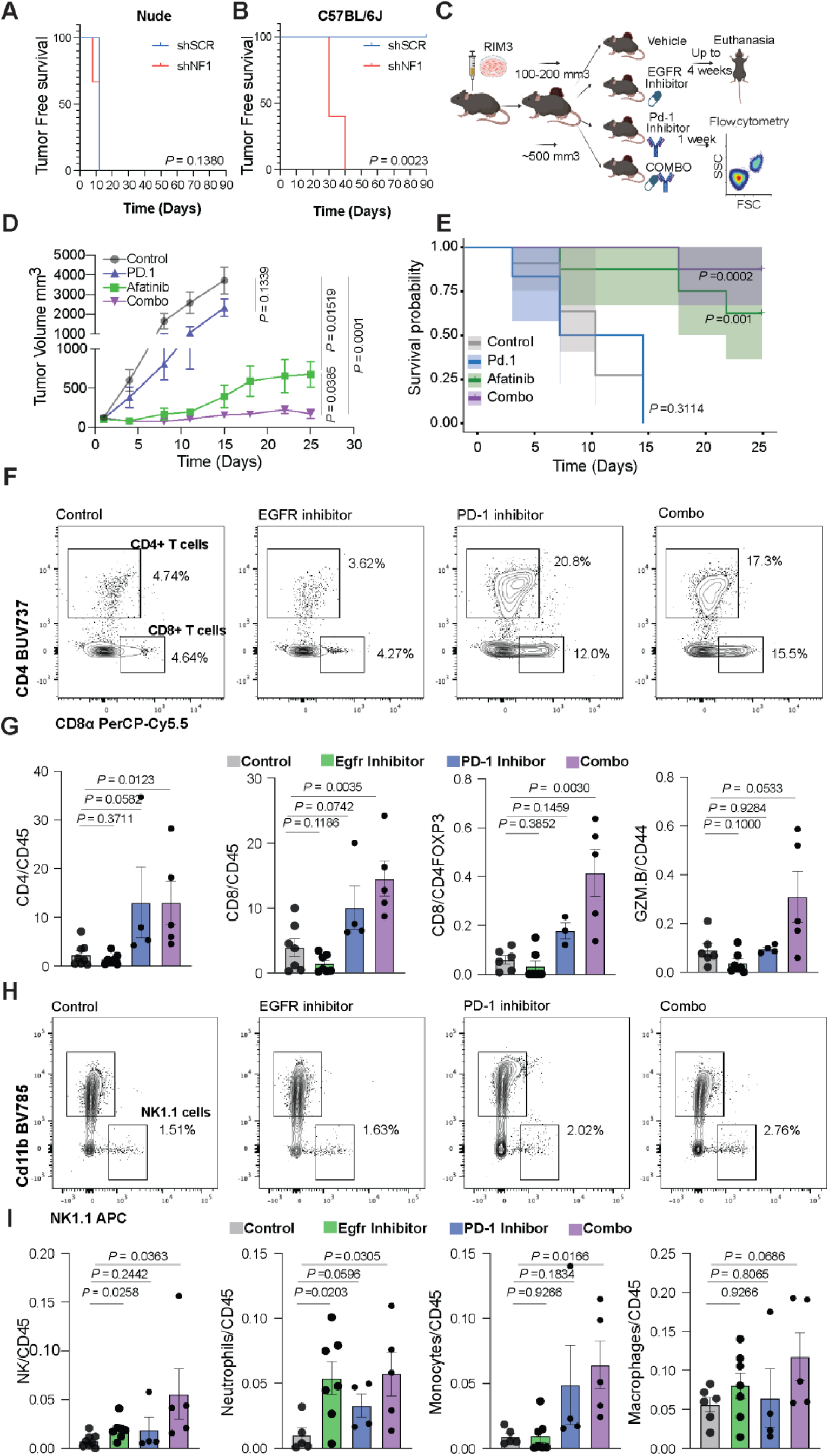
EGFR inhibition induces immune responses in *NF1* mutant melanoma preclinical models. **(A, B)** Kaplan Meier curves showing tumor-free survival of shNf1 or shSCR expressing RIM-3 cell lines in either immune-compromised nude (A) or immune-competent C57Bl/6J (B). P values calculated using log-rank test **(C)** Workflow of preclinical syngeneic model testing starting with transplanting shNf1 expression RIM-3 cells into BL/6J mice, monitoring tumor growth until either ∼100-200 mm^3^ for treatment or ∼ 500 mm^3^ for flow analysis, randomizing the mice into 4 treatment groups: control, EGFR inhibition, PD-1 inhibition or a combination of both, monitoring tumor growth and collecting the tumors at endpoint to generate single cell suspensions, staining with T or myeloid cells, and analyzing with FACS. **(D)** Growth curves of shNf1 RIM-3 melanoma cells treated with a vehicle control, afatinib, PD-1 mouse-specific antibody, or a combination. P-values were calculated using AUC. Data points represent mean ± SEM. **(E)** Kaplan Meier curves showing overall survival of (D). P values calculated using log-rank tests. **(F-I)** Flow gating, bar graphs, and data points of either T cells (F, G) or myeloid cells (H, I). Panels comparing mice after one week of treatment with control, afatinib, PD-1 antibody, or a combination of both. Bar graphs indicate mean ± SEM. P-values were calculated using a Mann-Whitney test.

To test whether these shNf1-expressing melanoma cells are resistant to immunotherapy and if EGFR inhibition can improve anti-PD-1 response, we randomized the mice into four groups that received afatinib, anti-PD-1 antibodies, afatinib and anti-PD-1 (combo), or a vehicle control **(Figure 7C)**. The anti-PD-1 antibody did not significantly inhibit tumor growth on its own, suggesting that shNf1-expressing RIM3 cells are ICI-resistant **(Figure 7D**). However, afatinib treatment inhibited tumor growth significantly compared to vehicle control, consistent with results from recently reported PDX models.^11^ A combination of afatinib with an anti-PD-1 antibody improved the tumor-inhibitory effect of afatinib even further in the shNf1-expressing RIM3 model. These responses resulted in prolonged survival of tumor-bearing mice treated with afatinib monotherapy or afatinib combined with anti-PD-1 until the experimental endpoint, whereas mice treated with vehicle control and anti-PD-1 monotherapy reached their humane endpoint sooner due to their rapid growth rates **(Figure 7E)**.

To determine how these therapies affect immune responses in these tumors, we treated an independent cohort of NF1-depleted RIM3 melanoma-bearing mice for one week and analyzed changes in myeloid and lymphoid cells using flow cytometry (FACS). We stained single cell suspensions with antibody panels that allowed us to detect and quantify different T cell (CD45, CD4, FOXP3, CD8, CD44, GZMB, kI67, TIM-3, and PD-1), or myeloid cell populations (CD45, NK-1.1, I-A/I-E, F4/80, CD11b, CD11c, Ly-6C, and Ly-6G). Our FACS data revealed that afatinib treatment alone did not affect tumor-infiltrating CD4 T cells or CD8 T cells. In contrast, PD-1 inhibition increased CD4 (p = 0.0582) and CD8 (p = 0.0742) T cell infiltration, although the response was variable and not statistically significant. In contrast, combined inhibition of EGFR and PD-1 led to a statistically significant increase in tumor-infiltrating CD4 (p = 0.0123) and CD8 (p = 0.0035) T cells. It also affected the CD8/Tregs ratio (p = 0.0030). Among T cell activity markers, GZMB increased (p = 0.053) in tumors treated with afatinib and anti-PD-1, but not in tumors treated with afatinib or anti-PD-1 alone (**Figure 7F, G**). These results indicate that combination therapy improves the T cell immune response, whereas anti-PD-1 or afatinib alone has no significant effect. Furthermore, afatinib treatment significantly increased neutrophil (p = 0.0203) and NK cell (p = 0.0258) infiltration in *Nf1* knockdown tumors (**Figure 7H, I**), perhaps due to upregulation of *Ulbp1* ^35^ **(Figure 6K)**. Anti-PD-1 antibodies had no significant effect on these cells, but effects on NK cells (p = 0.0363), neutrophils (p = 0.0305), and monocytes (p = 0.0166) were augmented by combination therapy (**Figure 7H, I**).

In sum, our data demonstrate that increased EGFR expression resulting from NF1 loss leads to immune evasion and, consequently, ICI resistance. Therefore, a combination of EGFR inhibition and ICI would be considered a more rational approach to improve clinical outcomes in patients with *NF1^Mut^* melanoma.

## Discussion

We deployed spatial transcriptomic and proteomic profiling of human melanomas, complemented by functional preclinical models, to define the multicellular architecture of the TME and to reveal its link to immune evasion and immunotherapy response. We identified 12 recurrent MNs as spatially correlated assemblies of transcriptionally distinct melanoma cells that consistently interact with specific stromal and immune populations across samples and patients. These MNs comprise melanoma cells in diverse states, together with subsets of CAFs, endothelial cells, T cells, myeloid populations, and other stromal cells, revealing modular spatial ecosystems that may underline clinical heterogeneity.

Given the unmet need for more effective treatment of *NF1^Mut^*melanoma,^9–12^ we focused on how NF1 loss shapes TME organization. We report that *NF1^Mut^* tumors were enriched for MNs marked by heightened EGF–EGFR signaling, NC-like dedifferentiation, more abundant CAFs, a collagen-rich extracellular matrix, and spatial patterns consistent with T cell exclusion. In contrast, *NF1^WT^*tumors exhibited greater infiltration by active, exhausted, and memory CD8 T cells. Our data also reveal that NF1 loss promotes EGFR expression and activation, thereby reducing MHC class I and II antigen presentation on melanoma cells and, in turn, promoting immune evasion. Consistent with these findings, we observed decreased HLA presentation in *NF1^Mut^* tumors and an inverse correlation between EGFR expression and intratumoral CD8 T cell density.

Our spatial analyses identified fewer MNs containing cytotoxic CD8 T cells, activated macrophages, and Tregs, as well as fewer intratumoral immune triads of CD4/CD8/DCs.^26^ Such an immune landscape is more consistent with impaired antigen presentation and T cell exclusion than with activation–exhaustion dynamics, in which exhausted and memory T cells would be expected to accumulate as more readily seen in *NF1^WT^* melanoma tissues. These results refine prior reports that emphasized PD-L1 upregulation as the dominant immune evasion mechanism in *NF1^Mut^* melanoma.^36^ Although increased PD-L1 surface expression has been observed in NF1-depleted cultures,^36^ and desmoplastic melanoma, a rare melanoma subtype that is typically driven by *NF1* mutations, has a high tumor mutational burden and responds favorably to immunotherapy,^3^ our spatial analyses of intact tissues indicate a more complex and immunologically “silent” phenotype in *NF1^Mut^* melanomas that is not adequately explained by checkpoint mediated inhibition alone.

CAFs may be central orchestrators of the *NF1^Mut^* TME. *NF1^Mut^* tumors exhibited a collagen-enriched matrix, increased CAF recruitment, and closer spatial proximity between CAFs and melanoma cells. Ligand–receptor analyses indicated significantly greater signaling from CAF populations to both tumor and neighboring stromal/immune cells in *NF1^Mut^*tumors, with associated pro-angiogenic programs. Together, these data suggest that NF1 loss remodels the extracellular matrix and amplifies CAF-driven signaling,^37^ creating an immune-suppressive, pro-angiogenic niche that physically and functionally restricts lymphocyte access to tumor cells.

Our results also expand on the broader literature implicating EGFR in tumor immune evasion in other solid tumors^16,38,39^, nominating this pathway as a potential target for combination therapy in *NF1^Mut^* melanoma. EGFR was enriched within MNs associated with increased oncogenic signaling activity and immune escape, and its expression correlated with immune-evasive transcriptional programs In *NF1^Mut^* melanoma. Functionally, we found that EGFR inhibition can restore antigen presentation in *NF1^Mut^*melanoma cells and enhance anti-tumor immunity. In addition to upregulating the MHC machinery,^40^ EGFR inhibition increased *ULBP1* expression, potentially facilitating NK cell recruitment and innate immune responses.^41^ These observations, together with the more permissive immune context in *NF1^WT^*tumors, support a therapeutic strategy that combines EGFR inhibition with PD-1 inhibitors to overcome antigen presentation and immune cell exclusion, thereby improving therapeutic responses in patients with *NF1^Mut^* melanoma.^17^ Results from our syngeneic mouse model support the feasibility and enhanced efficacy of this combinatorial treatment approach.

Our work has immediate implications for clinical translation. First, it delineates genotype-specific spatial ecosystems in melanoma linking NF1 loss to an EGFR-dependent immune-evasive program characterized by diminished antigen presentation, CAF-driven matrix remodeling, and T cell exclusion. Second, it nominates EGFR inhibition as a rational partner for PD-1 blockade in *NF1^Mut^* disease, providing a mechanistic basis for combination therapy. Third, it highlights spatially resolved biomarkers, including EGFR activity, HLA antigen presentation signatures, CAF density and proximity, collagen enrichment, and T cell exclusion metrics, that could guide patient stratification and pharmacodynamic monitoring.

We acknowledge some limitations. The Xenium targeted panel covers only ∼5,000 genes, so some signaling pathways may be missed. Although our dataset is the largest spatial multi-omic comparison of *NF1^Mut^* and *NF1^WT^* tissues, larger patient cohorts will need to be analyzed to assess the effects of variant-specific *NF1* mutations.

In conclusion, NF1 loss defines spatially distinct meta-niches in melanoma characterized by reduced MHC antigen presentation, CAF enrichment, matrix remodeling, and T cell exclusion, collectively associated with immune evasion. By linking these features to EGFR-dependent programs and demonstrating that EGFR inhibition can restore tumor immunogenicity, our findings provide a rationale for biomarker-driven clinical trials that combine EGFR inhibitors with PD-1 blockade in *NF1^Mut^* melanoma, or for adding EGFR inhibitors if *NF1^Mut^* melanomas do not respond to PD-1 monotherapy.

## Methods

### Study Cohort

Formalin-fixed paraffin-embedded (FFPE) tumor samples were prospectively collected from melanoma patients enrolled in the Interdisciplinary Melanoma Cooperative Group Database (IMCG) at NYU Langone Health (IRB #10362). Inclusion criteria for this clinicopathological database included patients who presented to the Laura and Isaac Perlmutter Cancer Center (2002-2018) with melanoma and were accrued within two months of diagnosis. All patients signed NYU Langone Health Institutional Review Board-approved consent authorizing the use of their specimens for research and follow-up.

### Xenium spatial analysis

#### Processing, Integration, and Annotation of Xenium Spatial Transcriptomics Data

Xenium analyses were performed on 42 FFPE tissues from 32 patients on two tissue microarrays. Data from multiple cores were acquired on the same Xenium Slide and imported into Seurat (v5.0.0)^42^ running on R (v4.3.2). Tissue coordinates of individual cores were imported as CSV files using the Xenium Explorer. For each core, a segmentation object was generated using CreateSegmentation(), and the core-specific field of view (FOV) was overlaid onto the full region FOV using Overlay(). Core-specific counts were extracted to create Seurat objects, filtering out cells with zero or low transcript counts (<50). Spatial features were reconstructed using centroid-based FOVs (CreateCentroids() and CreateFOV() and molecule overlays (CreateMolecules() for visualization within core boundaries. Individual core Seurat objects were first merged using the merge() function, with orig.ident retained to track the core of origin. The merged dataset was normalized and variance-stabilized using SCTransform() (selecting the top 5,000 variable features). Batch effects across patients were corrected using Harmony^43^ (HarmonyMatrix), with core ID specified as the grouping variable. Harmony-corrected embeddings were then used for downstream dimensionality reduction, including principal component analysis (PCA) and UMAP, as well as neighbor graph construction and clustering. Cell identities were assigned by identifying cluster-specific marker genes using FindAllMarkers() and manually annotating clusters based on canonical marker expression. Differential abundance of clusters and cell types between *NF1^Mut^* and *NF1^WT^* samples was calculated as the fraction of cells per patient in each cluster, and differences were assessed using Wilcoxon rank-sum tests with stat_compare_means() ggpubr (v0.6.2).^44^

#### Spatial Neighborhood Mapping and Meta-Niche Characterization

To characterize local cellular neighborhoods, tissue coordinates for each cell were retrieved, and, the composition of neighboring cell types within a 40 µm radius was computed for each cell using a fixed-radius nearest neighbor search (frNN function, dbscan package).^20^ Cells with fewer than three neighbors were excluded. The resulting neighborhood compositions were represented as vectors of cell-type fractions and clustered across all cells to identify 12 MNs using k-means clustering.^21^ MN labels were mapped back to the original Seurat objects, allowing for per-patient and per-cell assignment. Frequencies of MNs per patient were calculated, and differences between *NF1^Mut^* and *NF1^WT^* samples were assessed using Wilcoxon rank-sum tests. Niches that varied significantly between *NF1^Mut^* and *NF1^WT^*melanoma tissues were visualized using the ImageDimPlot function of Seurat with custom color schemes.

CellChat^24^ was used to infer cell-cell communication within spatially defined MNs or on each individual tissue core. Cells were grouped by cell type and *NF1* genotype, and interactions were predicted using the Human ligand–receptor database. Comparative network analyses identified shared, unique, and differential signaling between *NF1^Mut^*and *NF1^WT^* cells, with circle plots and heatmaps summarizing interaction strength and signaling roles across cell types.

#### Spatial proximity analysis of cell types

To quantify spatial relationships between cell types, tissue coordinates were extracted from each Seurat object, and the average nearest-neighbor distance between pairs of cell types within each core was computed using the get.knnx() function from the FNN package (v1.1.3). Cells with fewer than three neighbors were excluded. Distances were aggregated across cores by *NF1* genotype (mutant vs wild-type) to generate mean distance matrices. Fold-change matrices comparing *NF1^Mut^* to *NF1^WT^* distances were calculated and visualized using pheatmap (v1.0.12). Matrices of per-genotype distances, fold changes, and scaled values were exported for downstream interpretation.

#### Transcriptional profiling of spatial meta-niches and association with EGFR

Following MN identification, transcriptional programs were mapped by aggregating gene expression across cells assigned to each spatial MN. Average expression profiles were computed and used to characterize niche-specific transcriptional signatures. EGFR expression was quantified across meta-niches and summarized at the patient–niche level. Genotype-dependent differences between *NF1^Mut^* and *NF1^WT^*samples were assessed using Wilcoxon rank-sum tests. To resolve MN-specific mechanisms, both the fraction of EGFR-positive cells and the mean EGFR expression among EGFR-positive cells were evaluated, including stratification by dominant cell types within each MN. Pathway activity scores derived from Gene Ontology and Hallmark gene sets were correlated with EGFR expression using Spearman correlation analyses, thereby linking niche-specific transcriptional programs to EGFR signaling.

#### Cell type–specific differential expression and pathway enrichment independent of spatial niches

Differential expression analyses were performed on melanoma cells, CD8 T cells, CAFs, and myeloid cells independently of spatial niche assignment using Seurat. Cells were subsetted by cell type and stratified by *NF1* genotype. The RNA assay was normalized using NormalizeData(), and differential expression between *NF1^Mut^* and *NF1^WT^* cells was assessed using Wilcoxon rank-sum tests implemented in FindMarkers(). Genes with an adjusted *P*-value < 0.05 were considered significant and were separated into genotype-specific up- and downregulated sets. Functional enrichment analyses were conducted using clusterProfiler (v4.10.1) with gene identifier mapping via org.Hs.eg.db (v3.18.0). Gene Ontology Biological Process enrichment was performed using enrichGO(), and pathway-level enrichment was evaluated by ranking genes by log fold change, followed by GSEA using gseGO(). In parallel, Hallmark pathway enrichment was performed using Hallmark gene sets from the Molecular Signatures Database (MSigDB; collection H, release 2025.1), accessed via msigdbr (v25.1.1). Core enrichment genes were converted to gene symbols to facilitate biological interpretation. Data were visualized using ggplot2.^45^

### Bulk RNA-Seq

RNA-Seq data were analyzed as previously described.^11^ Briefly, Total RNA was extracted and treated with DNase using the RNeasy Plus Micro Kit (Qiagen, Cat#74034) following the manufacturer’s instructions. Libraries were prepared with the Illumina TruSeq Stranded Total RNA Ribo-Zero H/M/R Gold kit and sequenced on an Illumina NovaSeq 6000 at the NYU Genome Technology Center (RRID: SCR_017929). Sequencing data were demultiplexed and converted to FASTQ files using Illumina bcl2fastq. Reads were aligned to the human genome (hg19/GRCh37) or mouse genome (mm10) with the splice-aware STAR aligner, and PCR duplicates were removed using Picard (http://broadinstitute.github.io/picard/). Gene-level counts were generated with HTSeq, and DESeq2 was used to normalize counts and identify differentially expressed genes via negative binomial generalized linear models. Alignment, differential expression, and pathway analyses were performed using the Seq-N-Slide pipeline (RRID: SCR_021752, https://igordot.github.io/sns/)

### CODEX spatial proteomics

45-plex immunofluorescent (IF) imaging was performed on 56 archived FFPE tissues isolated from 32 melanoma patients. This 45-plex IF panel includes markers from Akoya Biosciences Phenocycler panel to classify melanoma cells (SOX10, S100b, GP100, NGFR, CSPG4, b-catenin, Axl), immune cells (CD20, CD3e, CD8, CD4, CD57, CD68, CD14, CD11c, CLEC9a, CD1c, Mac2, CD103, Foxp3, CD45RO, TCF-1, CD21, TOX, tryptase, CD169, CD66), endothelial cells (CD31, PDPN, LYVE1, AQP1, PNAd), epithelial cells (pan-cytokeratin) and stromal cells (PDPN, SMA, Collagen IV). This panel also includes markers for proliferation (Ki67), activation (IL-33, Granzyme B, IFNɣ), antigen presentation (HLA-DR), and immune inhibition (PD-1, ICOS, PD-L1, IDO1). 7 μm FFPE tissue was stained and scanned using PhenoCycler-Fusion 2.0 (Akoya Biosciences) according to Akoya’s commercial protocol. HALO Image Analysis Platform (Indica Labs, Albuquerque, NM USA), an AI assisted image segmentation software, was used for quantitative image analysis. We trained the software to accurately perform nuclear segmentation and identify tumor mass. This tumor classifier is trained by manually annotating tumor and non-tumor regions. The software considers both the structure and morphology of each pixel as well as the intensity of our selected markers (DAPI, SOX10, S100b, GP100, and CSPG4) to automatically segment cell borders and annotate tumor regions. Cells within tumor nests (i.e. CD8 T cells) are then annotated using markers threshold defined by pixel intensity and percent positive pixels within the cell borders. The final exported matrix contains the cell ID, tumor/non-tumor classification, average marker intensities, and the annotated cell phenotype. From this matrix, the cell densities (cells/mm2) and cell frequencies (cell count/total cells) are calculated for downstream analyses. An unpaired nonparametric Mann-Whitney test was performed to calculate p-values.

### Cell Culture

A375 (RRID: CVCL_0132), 08-175,^11^ SK-MEL-28 (RRID:CVCL_0526) and RIM3^33^ cell lines were utilized in this study.^46^ A375 and Sk-MEL-28 cells were cultured in Corning™ RPMI 1640 Medium (Mod.) 1X with L-Glutamine (Life Sciences, 10-040-CV) and RIM-3 cells were cultured in DMEM (Corning, 10-017-CV). All culture media were enriched with 10% fetal bovine serum (FBS), L-glutamine from Invitrogen, sodium pyruvate solution (Sigma, S8636), non-essential amino acids (Sigma, M7145), GlutaMAX supplement (Gibco, 25050061) and normocin (Invivogen, NC9273499). All cell lines were maintained at 37°C in 7% CO2 and were periodically tested for mycoplasma. Human cell lines *NF1* and *EGFR* knockdowns were previously described.^11^ All Lentiviral and piggyback vectors were designed and purchased from Vector Builder (Chicago, USA). shNF1 target sequence: GCCAACCTTAACCTCTCTAAT, and shEGFR target sequence: GCATAGGCATTGGTGAATTTA. NF1 ectopic expression was performed by using a piggyback vector that expresses a flag-tagged, tetracycline-inducible NF1 full-length gene (Vector builder: VB250317-1543ver), and an empty flag (VB250319-1011vbt) was used as a control. Plasmids were transfected using Lipofectamine 3000 (Invitrogen, L3000015) and expression was induced by adding 1ug/ml doxycycline (Gold-Bio, D-500-1) for 48 hours. Western blot and qPCR were performed as previously described^11^ using NF1 Abcam, ab17963 (RRID: AB_444142)), EGFR (Abcam, ab52894 (RRID: AB_869579)), HSP90 (Cell Signaling, 4877 (RRID: AB_2233307)), HLA class I ABC Polyclonal Antibody (ThermoFisher, 15240-1-AP (AB_1557426)) and horseradish peroxidase-conjugated secondary antibodies (Cell Signaling 7074 (RRID: AB_2099233) and 7076 (RRID: AB_330924)). qPCR primers were ordered from Invitrogen, USA. Primers used were: NF1: F: AACTTCTTCCTGCGACTGCG, R: CTCTGCGACAGACGTCAACA. EGFR: F: ACCTCTCCCGGTCAGAGATG, R: TGTGCCTTGGCAGACTTTCT. Live cell imaging was performed using Incucyte (Essen BioScience) as described before^11^. Flow cytometry analysis was performed by staining cells treated with Human IFN-gamma Recombinant Protein (Thermo-Fischer, Cat # 300-02-100U) with PE anti-human HLA-A,B,C (Biolegend 311406 (RRID: AB_314875) or APC anti-human HLA-DR, DP, DQ Antibody (Biolegend 361714 (RRID: AB_2750316)) at 4 °C for 30 minutes in the dark. Flow analysis was performed using BD FACS Symphony A5 analyzer and analyzed using FlowJo V10.10.0.

### Mouse Models

RIM3 melanoma cell lines (expressing shSCR or shNf1) were used for subcutaneous flank injection. For tumor growth assays, 1 × 10⁶ cells were transplanted, whereas 2 × 10⁵ cells were used for treatment experiments. Cells were injected in six-week-old male nude mice (Jackson Laboratories, RRID:IMSR_JAX:002019) or six-week-old male C57BL/6J mice (Jackson Laboratories, RRID:IMSR_JAX:000664). Tumor growth was monitored and measured with digital calipers (in millimeters). Tumor volumes were calculated with the formula: (v (π/6*l*w2), where l = length in mm, w = width in mm, v=mm3). shNF1 RIM-3 tumors were allowed to reach ∼100-200 mm^3^ for growth curves or ∼500 mm^3^ for flow analysis before being randomly assigned to either afatinib treatment (Medchem Express, BIBW 2992, Cat# HY-10261), PD-1(CD279) anti-mouse Mab (Bioxcell, Cat# BE0146), a combination of both treatments or the appropriate vehicle control. Afatinib was administrated at 20mg/kg five days a week by gastric gavage. PD-1 Mab was administrated twice weekly at a concentration of 10 mg/kg by intraperitoneal injection. Treatments were administrated for up to four weeks for growth curves or one week for flow analysis. To stain for immune markers, tumor masses were measured at the endpoint. Resected tumors were digested at 37 °C for 25 minutes with agitation by Collagenase D (1 mg/mL): (Sigma-Aldrich Cat#11088866001) and DNase I (80 U/mL) (Roche Cat#04536282001) and passed through 70 uM cell strainers to generate single cell suspensions. Each tumor was divided into two wells and stained with a T cell panel or a Myeloid panel in addition to Ghost Dye live/dead stain (CYTEK Cat#130865T100). T cell panel included: CD45 (BD Biosciences Cat#564279 (RRID: AB_2651134)), CD4 (BD Biosciences Cat#612844), CD8a (CYTEK, Cat#6 5-0081-U100), CD44 (CYTEK, Cat#60-0441-U100), GZMB (ThermoFisher Cat# MHGB04 (RRID: AB_10372671)), Ki67 (ThermoFisher Cat#48-5698-80 (RRID: AB_11151155)), FOXP3 (BioLegend Cat#126407 (RRID: AB_1089116)), PD-1 (BioLegened Cat#135225 (RRID: AB_2563680)), )), TIM-3 (BioLegened Cat#75833-230) anti-mouse antibodies. Myeloid panel included:, NK1.1 (Biolegend Cat# 108710 (RRID: AB_313397)), CD11b (Biolegend Cat#101243 (RRID: AB_2561373)), CD11c (Biolegend Cat#117333 (RRID: AB_11204262)), Ly-6C (Biolegend Cat# 128012 (RRID: AB_1659241)), Ly-6G (Biolegend Cat#127618 (RRID: AB_1877261)), I-A/I-E (Biolegend Cat# 107632 (RRID: AB_2650896)), F4/80 (ThermoFisher Cat#12-4801-82 (RRID: AB_465923)) anti-mouse antibodies. Tumors were stained for 1 hour at 4°C in the dark. After washing, tumors were fixed in 4% paraformaldehyde and stored at 4°C until analysis. Flow analysis was performed using BD FACS Symphony A5 analyzer and analyzed using FlowJo V10.10.0. Number of cells were normalized to tumor mass and Mann Whitney test was used to compare groups.

### Statistical Analysis

Statistical analyses were performed with Microsoft Excel, GraphPad Prism (version 10.2.0, RRID:SCR_002798), or R statistical software (RRID:SCR_001905 version 4.3.1).

## Data Availability statement

All spatial and RNA-Seq will be available in public domains (GEO).

## Acknowledgements

Xenium and Codex sequencing were performed at the Experimental Pathology Core. RNA-Seq library preps and sequencing were performed at the Genome Technology Center at NYU Grossman School of Medicine.

The authors are grateful to Dr. Lukas Sommer from the University of Zurich for providing the RIM-3 melanoma cells used in this study. We are also grateful to Drs. Matthew Klairmont and Christopher Park for discussions and advice on Xenium data handling and analysis.

## Author contributions

**Project concept and design:** M. Ibrahim, Amanda W. Lund, M. Schober, and I. Osman.

**Clinical evaluation, pathology evaluation, and selection of the melanoma patients for the experiments:** G. Jour and I. Osman.

**Experiments execution and acquisition of data:** M. Ibrahim, I. Illa-Bochaca, Tara Muijlwijk, Ines Delclaux, Katherine S Ventre, Paola Angulo Salgado Shi Qiu and Agrima Dutt

**Data analysis and interpretation, figure preparation:** M. Ibrahim, Tara Muijlwijk, Ines Delclaux, Katherine S Ventre, Paola Angulo Salgado, Amanda W. Lund, M. Schober, and I. Osman.

**Writing, reviewing, and/or revising the manuscript:** M. Ibrahim, M. Schober, and I. Osman. All authors revised the manuscript and gave final approval for publication.

---

**Figure S1.**
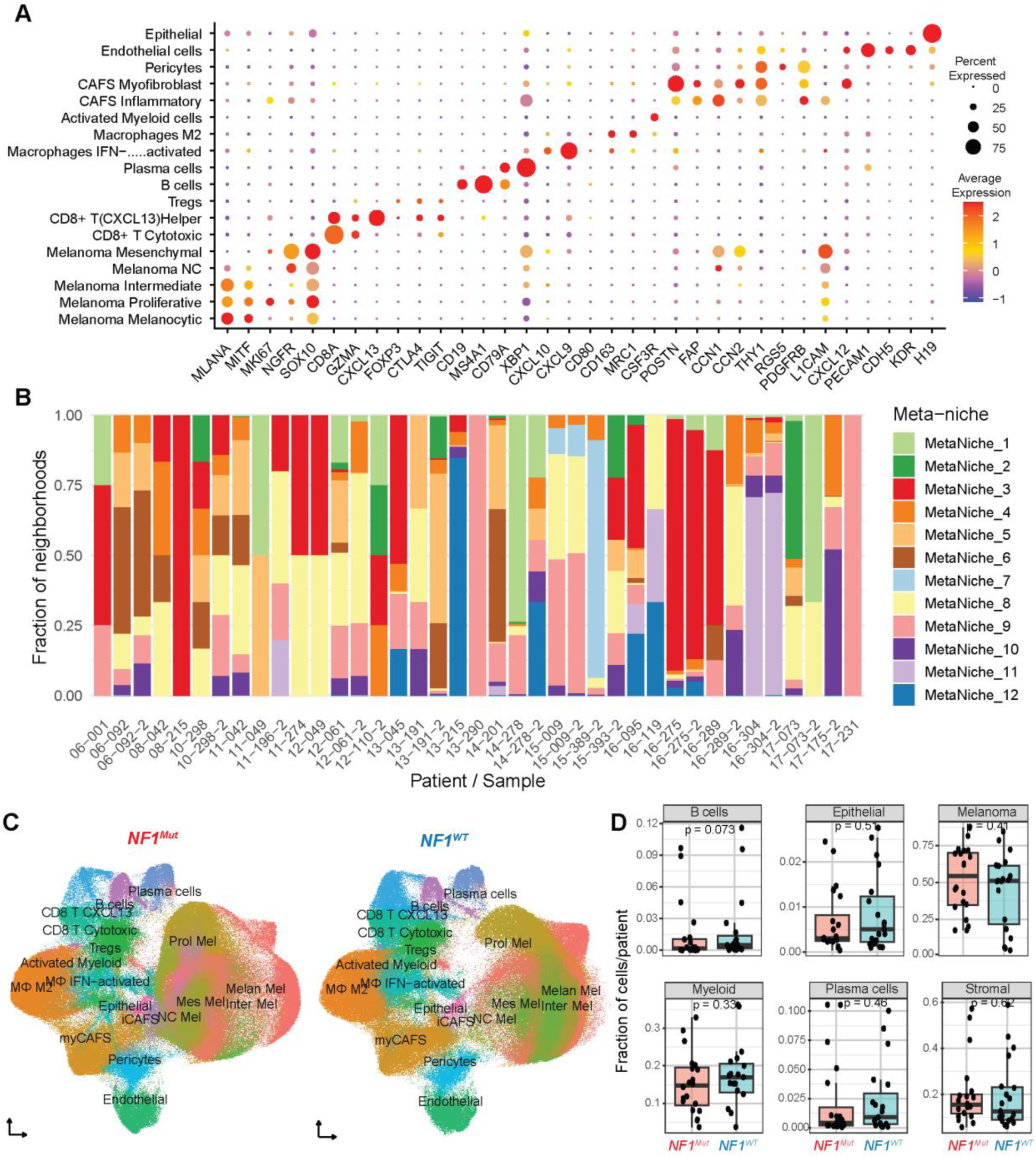
The composition of cell clusters and meta-niches in melanoma. **(A)** Dot plot showing canonical marker genes for each cell type identified in Xenium analysis. **(B)** Stacked bar plot showing the ratio of each meta-niche identified in each sample. **(C)** UMAP plots of lineage sub-types identified in *NF1^Mut^* Vs. *NF^WT^*samples. **(D)** Box plots and data points of the proportion of major cell types identified in *NF1^Mut^*and *NF^WT^* melanoma patients. P-values indicate a Wilcox test.

**Figure S2.**
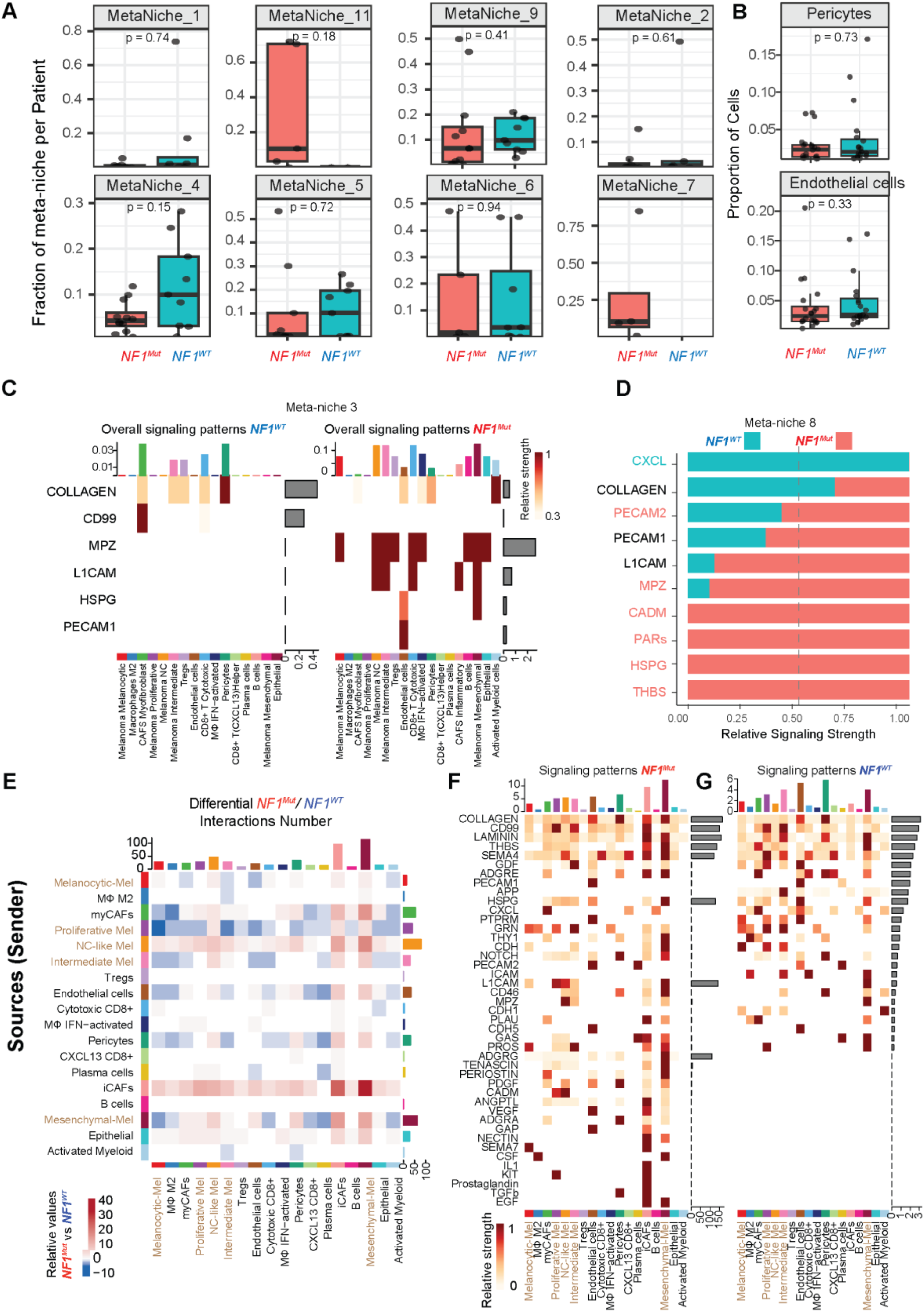
Cell-Cell interactions promote immune evasion in *NF1* Mutant melanoma microenvironment. **(A)** Box plots and data points of the proportion of meta-niches per number of cells in *NF1^Mut^* and *NF^WT^* melanoma patients. P-values indicate a Wilcox test. **(B)** Box plots and data points of the proportion of pericytes and endothelial cells out of total cells in *NF1^Mut^* and *NF^WT^* melanoma patients. P-values indicate a Wilcox test. **(C)** Heatmaps showing signaling patterns in *NF1^Mut^* meta-niche 3. **(D)** CellChat Bar plot showing cell-cell signaling in *NF^WT^* vs. *NF1^Mut^* meta-niche 8. The teal label indicates signaling is significantly higher in *NF^WT^*, the red label indicates signaling is significantly higher in *NF1^Mut^*, and the black label indicates no significant difference. **(E)** CellChat heatmaps showing *the* number *of* differential interactions between *NF1^Mut^* and *NF^WT^*. Senders are in rows and receivers are in columns. Red color indicates more, while blue color indicates fewer interactions in *NF1^Mut^*. **(F, G)** CellChat heatmaps showing signaling patterns in *NF1^Mut^* (F) or *NF^WT^*(G) microenvironment. The darker the color, the stronger the signaling strength.

**Figure S3.**
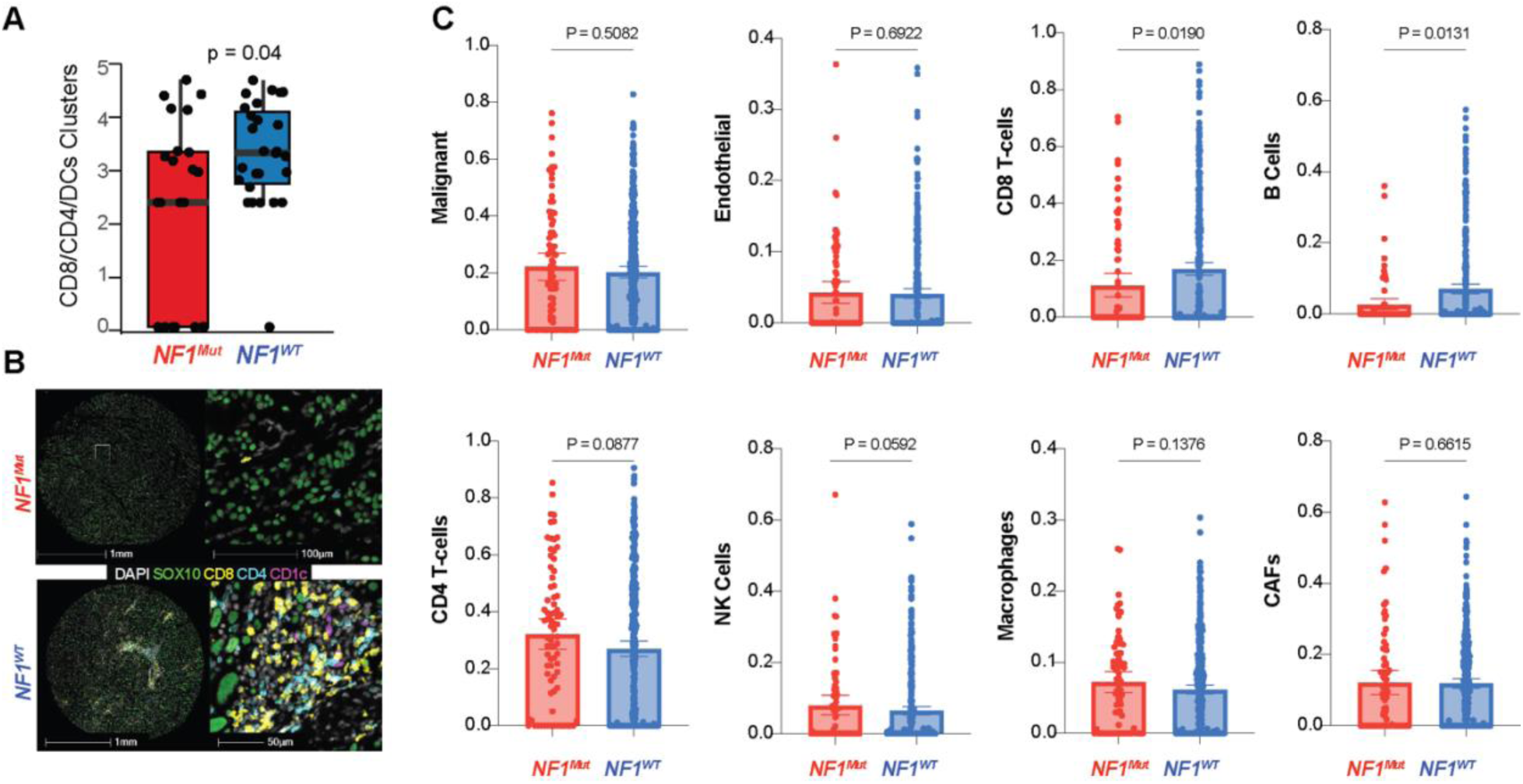
*NF1* mutant melanoma cells are characterized by fewer infiltrating CD8 T cells. **(A)** Box plots and data points of proximity scores of CD4/CD8/DCs immune clusters in *NF1^Mut^* and *NF^WT^* tissue. P-values indicate a Wilcox test. (**B**) Representative images of melanoma tissues showing clusters of CD4, CD8, and CD1c in *NF1^Mut^* or *NF^WT^***. (C)** Box plots and data points of CIBERSORTx analysis showing different cell clusters identified in the TCGA SCKM dataset. P-values were calculated using a Mann-Whitney test.

**Figure S4.**
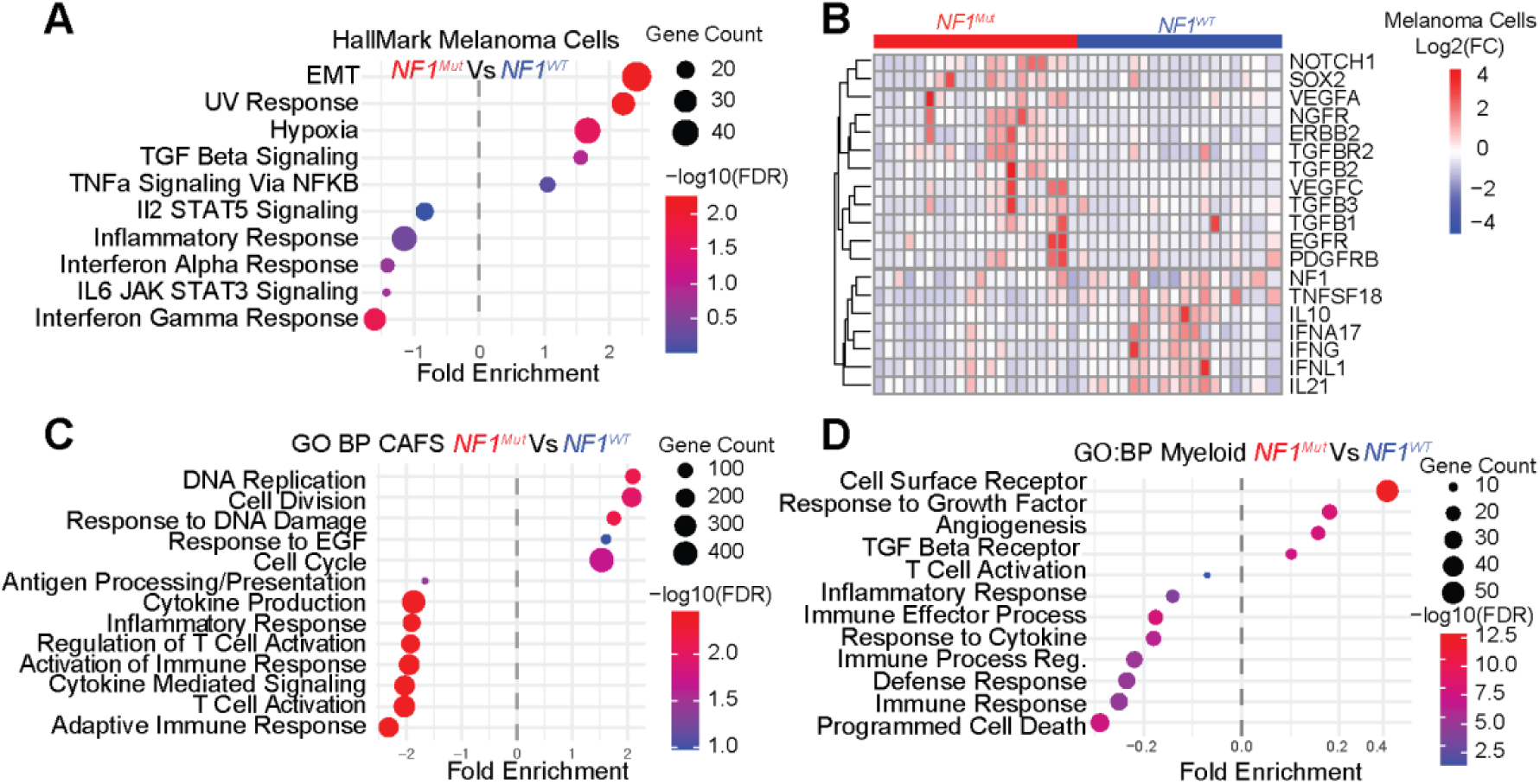
EGFR signaling is enriched in tumor, stromal, and immune cells in *NF1* mutant melanoma. **(A)** GSEA Hallmark pathway analysis showing EMT, TGF beta, and hypoxia are upregulated, and interferon alpha and gamma and inflammatory response are downregulated in melanoma clusters of *NF1^Mut^* tumors. **(B)** Heatmap of genes significantly differentially expressed between *NF1^Mut^* and *NF^WT^*melanoma clusters. **(C)** GO pathway analysis shows that DNA replication and response to EGFR are upregulated, while T cell activation, adaptive immune response, and cytokine production are downregulated in CAF clusters of *NF1^Mut^* melanoma. **(D)** GO pathway analysis shows that response to growth factor and TGFβ signaling are upregulated, and immune response, inflammatory response, and programmed cell death are downregulated in myeloid (macrophages, M2, IFN-γ–activated, and activated myeloid cells) clusters of *NF1^Mut^* melanoma.

## References

1. Shain, A.H., Yeh, I., Kovalyshyn, I., Sriharan, A., Talevich, E., Gagnon, A., Dummer, R., North, J., Pincus, L., Ruben, B., et al. (2015). The Genetic Evolution of Melanoma from Precursor Lesions. New England Journal of Medicine 373, 1926–1936. doi:10.1056/NEJMoa1502583.

2. Schiantarelli, J., Benamar, M., Park, J., Sax, H.E., Oliveira, G., Bosma-Moody, A., Campbell, K.M., Liu, D., Johnson, D.B., Rodig, S., et al. (2025). Genomic mediators of acquired resistance to immunotherapy in metastatic melanoma. Cancer Cell 43, 308–316.e306. 10.1016/j.ccell.2025.01.009.

3. Eroglu, Z., Zaretsky, J.M., Hu-Lieskovan, S., Kim, D.W., Algazi, A., Johnson, D.B., Liniker, E., Ben, K., Munhoz, R., Rapisuwon, S., et al. (2018). High response rate to PD-1 blockade in desmoplastic melanomas. Nature 553, 347–350. 10.1038/nature25187.

4. Andor, N., Graham, T.A., Jansen, M., Xia, L.C., Aktipis, C.A., Petritsch, C., Ji, H.P., and Maley, C.C. (2016). Pan-cancer analysis of the extent and consequences of intratumor heterogeneity. Nat Med 22, 105–113. 10.1038/nm.3984.

5. Tirosh, I., and Suva, M.L. (2024). Cancer cell states: Lessons from ten years of single-cell RNA-sequencing of human tumors. Cancer Cell 42, 1497–1506. 10.1016/j.ccell.2024.08.005.

6. Curti, B.D., and Faries, M.B. (2021). Recent Advances in the Treatment of Melanoma. New England Journal of Medicine 384, 2229–2240. doi:10.1056/NEJMra2034861.

7. Tasdogan, A., Sullivan, R.J., Katalinic, A., Lebbe, C., Whitaker, D., Puig, S., van de Poll-Franse, L.V., Massi, D., and Schadendorf, D. (2025). Cutaneous melanoma. Nature Reviews Disease Primers 11, 23. 10.1038/s41572-025-00603-8.

8. Tirosh, I., Izar, B., Prakadan, S.M., Wadsworth, M.H., 2nd, Treacy, D., Trombetta, J.J., Rotem, A., Rodman, C., Lian, C., Murphy, G., et al. (2016). Dissecting the multicellular ecosystem of metastatic melanoma by single-cell RNA-seq. Science 352, 189–196. 10.1126/science.aad0501.

9. Jour, G., Illa-Bochaca, I., Ibrahim, M., Donnelly, D., Zhu, K., Miera, E.V., Vasudevaraja, V., Mezzano, V., Ramswami, S., Yeh, Y.H., et al. (2023). Genomic and Transcriptomic Analyses of NF1-Mutant Melanoma Identify Potential Targeted Approach for Treatment. J Invest Dermatol 143, 444–455.e448. 10.1016/j.jid.2022.07.022.

10. Cirenajwis, H., Lauss, M., Ekedahl, H., Torngren, T., Kvist, A., Saal, L.H., Olsson, H., Staaf, J., Carneiro, A., Ingvar, C., et al. (2017). NF1-mutated melanoma tumors harbor distinct clinical and biological characteristics. Mol Oncol 11, 438–451. 10.1002/1878-0261.12050.

11. Ibrahim, M., Illa-Bochaca, I., Jour, G., Vega-Saenz de Miera, E., Fracasso, J., Ruggles, K., Osman, I., and Schober, M. (2025). NF1 Loss Promotes EGFR Activation and Confers Sensitivity to EGFR Inhibition in NF1-Mutant Melanoma. Cancer Res 85, 3348–3364. 10.1158/0008-5472.Can-24-3904.

12. Nissan, M.H., Pratilas, C.A., Jones, A.M., Ramirez, R., Won, H., Liu, C., Tiwari, S., Kong, L., Hanrahan, A.J., Yao, Z., et al. (2014). Loss of NF1 in cutaneous melanoma is associated with RAS activation and MEK dependence. Cancer Res 74, 2340–2350. 10.1158/0008-5472.Can-13-2625.

13. Maertens, O., Johnson, B., Hollstein, P., Frederick, D.T., Cooper, Z.A., Messiaen, L., Bronson, R.T., McMahon, M., Granter, S., Flaherty, K., et al. (2013). Elucidating distinct roles for NF1 in melanomagenesis. Cancer Discov 3, 338–349. 10.1158/2159-8290.Cd-12-0313.

14. Cancer Genome Atlas, N. (2015). Genomic Classification of Cutaneous Melanoma. Cell 161, 1681–1696. 10.1016/j.cell.2015.05.044.

15. Thielmann, C.M., Chorti, E., Matull, J., Murali, R., Zaremba, A., Lodde, G., Jansen, P., Richter, L., Kretz, J., Möller, I., et al. (2021). NF1-mutated melanomas reveal distinct clinical characteristics depending on tumour origin and respond favourably to immune checkpoint inhibitors. Eur J Cancer 159, 113–124. 10.1016/j.ejca.2021.09.035.

16. Fasano, M., Della Corte, C.M., Viscardi, G., Di Liello, R., Paragliola, F., Sparano, F., Iacovino, M.L., Castrichino, A., Doria, F., Sica, A., et al. (2021). Head and neck cancer: the role of anti-EGFR agents in the era of immunotherapy. Ther Adv Med Oncol 13, 1758835920949418. 10.1177/1758835920949418.

17. Sacco, A.G., Chen, R., Worden, F.P., Wong, D.J.L., Adkins, D., Swiecicki, P., Chai-Ho, W., Oppelt, P., Ghosh, D., Bykowski, J., et al. (2021). Pembrolizumab plus cetuximab in patients with recurrent or metastatic head and neck squamous cell carcinoma: an open-label, multi-arm, non-randomised, multicentre, phase 2 trial. Lancet Oncol 22, 883–892. 10.1016/s1470-2045(21)00136-4.

18. Madeddu, C., Donisi, C., Liscia, N., Lai, E., Scartozzi, M., and Macciò, A. (2022). EGFR-Mutated Non-Small Cell Lung Cancer and Resistance to Immunotherapy: Role of the Tumor Microenvironment. Int J Mol Sci 23. 10.3390/ijms23126489.

19. Tsoi, J., Robert, L., Paraiso, K., Galvan, C., Sheu, K.M., Lay, J., Wong, D.J.L., Atefi, M., Shirazi, R., Wang, X., et al. (2018). Multi-stage Differentiation Defines Melanoma Subtypes with Differential Vulnerability to Drug-Induced Iron-Dependent Oxidative Stress. Cancer Cell 33, 890–904.e895. 10.1016/j.ccell.2018.03.017.

20. Chen, X., and Güttel, S. (2024). Fast and exact fixed-radius neighbor search based on sorting. PeerJ Comput Sci 10, e1929. 10.7717/peerj-cs.1929.

21. Hartigan, J.A., and Wong, M.A. (2018). A K-Means Clustering Algorithm. Journal of the Royal Statistical Society Series C: Applied Statistics 28, 100–108. 10.2307/2346830.

22. Hossain, S.M., Gimenez, G., Stockwell, P.A., Tsai, P., Print, C.G., Rys, J., Cybulska-Stopa, B., Ratajska, M., Harazin-Lechowska, A., Almomani, S., et al. (2022). Innate immune checkpoint inhibitor resistance is associated with melanoma sub-types exhibiting invasive and de-differentiated gene expression signatures. Front Immunol 13, 955063. 10.3389/fimmu.2022.955063.

23. Schaaf, M.B., Garg, A.D., and Agostinis, P. (2018). Defining the role of the tumor vasculature in antitumor immunity and immunotherapy. Cell Death & Disease 9, 115. 10.1038/s41419-017-0061-0.

24. Jin, S., Guerrero-Juarez, C.F., Zhang, L., Chang, I., Ramos, R., Kuan, C.-H., Myung, P., Plikus, M.V., and Nie, Q. (2021). Inference and analysis of cell-cell communication using CellChat. Nature Communications 12, 1088. 10.1038/s41467-021-21246-9.

25. Maten, M.V., Reijnen, C., Pijnenborg, J.M.A., and Zegers, M.M. (2019). L1 Cell Adhesion Molecule in Cancer, a Systematic Review on Domain-Specific Functions. Int J Mol Sci 20. 10.3390/ijms20174180.

26. Espinosa-Carrasco, G., Chiu, E., Scrivo, A., Zumbo, P., Dave, A., Betel, D., Kang, S.W., Jang, H.-J., Hellmann, M.D., Burt, B.M., et al. (2024). Intratumoral immune triads are required for immunotherapy-mediated elimination of solid tumors. Cancer Cell 42, 1202–1216.e1208. 10.1016/j.ccell.2024.05.025.

27. Newman, A.M., Liu, C.L., Green, M.R., Gentles, A.J., Feng, W., Xu, Y., Hoang, C.D., Diehn, M., and Alizadeh, A.A. (2015). Robust enumeration of cell subsets from tissue expression profiles. Nature Methods 12, 453–457. 10.1038/nmeth.3337.

28. Hazini, A., Fisher, K., and Seymour, L. (2021). Deregulation of HLA-I in cancer and its central importance for immunotherapy. J Immunother Cancer 9. 10.1136/jitc-2021-002899.

29. Hsieh, C.H., Jian, C.Z., Lin, L.I., Low, G.S., Ou, P.Y., Hsu, C., and Ou, D.L. (2022). Potential Role of CXCL13/CXCR5 Signaling in Immune Checkpoint Inhibitor Treatment in Cancer. Cancers (Basel) 14. 10.3390/cancers14020294.

30. Seitz, S., Dreyer, T.F., Stange, C., Steiger, K., Bräuer, R., Scheutz, L., Multhoff, G., Weichert, W., Kiechle, M., Magdolen, V., and Bronger, H. (2022). CXCL9 inhibits tumour growth and drives anti-PD-L1 therapy in ovarian cancer. British Journal of Cancer 126, 1470–1480. 10.1038/s41416-022-01763-0.

31. Oughtred, R., Rust, J., Chang, C., Breitkreutz, B.J., Stark, C., Willems, A., Boucher, L., Leung, G., Kolas, N., Zhang, F., et al. (2021). The BioGRID database: A comprehensive biomedical resource of curated protein, genetic, and chemical interactions. Protein Sci 30, 187–200. 10.1002/pro.3978.

32. Zingg, D., Debbache, J., Schaefer, S.M., Tuncer, E., Frommel, S.C., Cheng, P., Arenas-Ramirez, N., Haeusel, J., Zhang, Y., Bonalli, M., et al. (2015). The epigenetic modifier EZH2 controls melanoma growth and metastasis through silencing of distinct tumour suppressors. Nature Communications 6, 6051. 10.1038/ncomms7051.

33. Zingg, D., Arenas-Ramirez, N., Sahin, D., Rosalia, R.A., Antunes, A.T., Haeusel, J., Sommer, L., and Boyman, O. (2017). The Histone Methyltransferase Ezh2 Controls Mechanisms of Adaptive Resistance to Tumor Immunotherapy. Cell Reports 20, 854–867. 10.1016/j.celrep.2017.07.007.

34. Solca, F., Dahl, G., Zoephel, A., Bader, G., Sanderson, M., Klein, C., Kraemer, O., Himmelsbach, F., Haaksma, E., and Adolf, G.R. (2012). Target binding properties and cellular activity of afatinib (BIBW 2992), an irreversible ErbB family blocker. J Pharmacol Exp Ther 343, 342–350. 10.1124/jpet.112.197756.

35. Bauman, Y., Drayman, N., Ben-Nun-Shaul, O., Vitenstein, A., Yamin, R., Ophir, Y., Oppenheim, A., and Mandelboim, O. (2016). Downregulation of the stress-induced ligand ULBP1 following SV40 infection confers viral evasion from NK cell cytotoxicity. Oncotarget 7.

36. Berry, D., Moldoveanu, D., Rajkumar, S., Lajoie, M., Lin, T., Tchelougou, D., Sakthivel, S., Sharon, I., Bernard, A., Pelletier, S., et al. (2025). The NF1 tumor suppressor regulates PD-L1 and immune evasion in melanoma. Cell Rep 44, 115365. 10.1016/j.celrep.2025.115365.

37. Sahai, E., Astsaturov, I., Cukierman, E., DeNardo, D.G., Egeblad, M., Evans, R.M., Fearon, D., Greten, F.R., Hingorani, S.R., Hunter, T., et al. (2020). A framework for advancing our understanding of cancer-associated fibroblasts. Nature Reviews Cancer 20, 174–186. 10.1038/s41568-019-0238-1.

38. Santaniello, A., Napolitano, F., Servetto, A., De Placido, P., Silvestris, N., Bianco, C., Formisano, L., and Bianco, R. (2019). Tumour Microenvironment and Immune Evasion in EGFR Addicted NSCLC: Hurdles and Possibilities. Cancers (Basel) 11. 10.3390/cancers11101419.

39. Wang, X., Semba, T., Manyam, G.C., Wang, J., Shao, S., Bertucci, F., Finetti, P., Krishnamurthy, S., Phi, L.T.H., Pearson, T., et al. (2022). EGFR is a master switch between immunosuppressive and immunoactive tumor microenvironment in inflammatory breast cancer. Science Advances 8, eabn7983. doi:10.1126/sciadv.abn7983.

40. Passarelli, A., Mannavola, F., Stucci, L.S., Tucci, M., and Silvestris, F. (2017). Immune system and melanoma biology: a balance between immunosurveillance and immune escape. Oncotarget 8, 106132–106142. 10.18632/oncotarget.22190.

41. Kim, H., Kim, S.H., Kim, M.J., Kim, S.J., Park, S.J., Chung, J.S., Bae, J.H., and Kang, C.D. (2011). EGFR inhibitors enhanced the susceptibility to NK cell-mediated lysis of lung cancer cells. J Immunother 34, 372–381. 10.1097/CJI.0b013e31821b724a.

42. Hao, Y., Stuart, T., Kowalski, M.H., Choudhary, S., Hoffman, P., Hartman, A., Srivastava, A., Molla, G., Madad, S., Fernandez-Granda, C., and Satija, R. (2024). Dictionary learning for integrative, multimodal and scalable single-cell analysis. Nature Biotechnology 42, 293–304. 10.1038/s41587-023-01767-y.

43. Korsunsky, I., Millard, N., Fan, J., Slowikowski, K., Zhang, F., Wei, K., Baglaenko, Y., Brenner, M., Loh, P.-r., and Raychaudhuri, S. (2019). Fast, sensitive and accurate integration of single-cell data with Harmony. Nature Methods 16, 1289–1296. 10.1038/s41592-019-0619-0.

44. Kassambara, A. (2018). ggpubr:’ggplot2’based publication ready plots. R package version, 2.

45. Wickham, H. (2016). ggplot2: Elegant Graphics for Data Analysis (Springer International Publishing).

46. Ibrahim, M., Illa-Bochaca, I., Fa’ak, F., Monson, K.R., Ferguson, R., Lyu, C., Vega-Saenz de Miera, E., Johannet, P., Chou, M., Mastroianni, J., et al. (2023). Kinase Insert Domain Receptor Q472H Pathogenic Germline Variant Impacts Melanoma Tumor Growth and Patient Treatment Outcomes. Cancers (Basel) 16. 10.3390/cancers16010018.

